# CRISPR screening reveals genetic regulators associated with the evolution of eye degeneration

**DOI:** 10.64898/2025.12.30.697043

**Authors:** Devin Shennard, Charlotte Mulliniks, Estephany Ferrufino, Aubrey Manning, Stefan Choy, Rianna Ambosie, Shiyanth Thevasagayampillai, Brooke Pileggi, Lane Zimmermann, Chenxiao Shen, Alli Kimmel, Renee Mapa, Preethi H. Gunaratne, Erik R. Duboue, Wynn K. Meyer, Beau Alward, Suzanne McGaugh, Johanna E. Kowalko

**Affiliations:** Department of Biological Sciences, Lehigh University, Bethlehem, PA; Department of Biology and Biochemistry, University of Houston, Houston, TX; Human Genome Sequencing Center, Baylor College of Medicine, Houston, TX; Wilkes Honors College, Florida Atlantic University, Jupiter, FL; Department of Integrative Biology and Physiology, University of California, Los Angeles, Los Angeles, CA; Ecology, Evolution, and Behavior, 140 Gortner Lab, 1479 Gortner Ave, University of Minnesota, Saint Paul, MN 55108, USA

## Abstract

Determining the genetic factors contributing to trait evolution is critical for understanding how and why traits evolve; however, establishing which genes underlie the evolution of complex traits remains challenging. The freshwater fish *Astyanax mexicanus*, a species that includes blind, cave-dwelling and eyed, surface-dwelling fish, is a powerful model for evolutionary genetics. While genetic mapping studies in this species previously identified genomic regions associated with cave-derived traits, few causative genes and genetic changes have been identified. Here, we develop methods to identify and rapidly functionally assess candidate genes in *A. mexicanus*, focusing on a defining trait of cave animals, eye loss. Candidate genes were identified based on whether they fell within an eye-related quantitative trait locus, were differentially expressed between surface and cave eyes, and showed evidence of positive selection in cavefish. Single-nucleus RNA-sequencing revealed that these candidate genes were expressed in multiple cell types during development, including those in different tissues of the eye. CRISPR-Cas9-based mutagenesis demonstrated that disruption of nine of these candidate genes in surface fish resulted in altered eye size. Perturbation of one of these genes, *fibulin-7* (*fbln7*), revealed changes in eye size across multiple stages of eye development. Together, this work identified multiple genes associated with the evolution of eye degeneration in *A. mexicanus.* Further, this study represents a roadmap for rapid identification and functional assessment of candidate genes implicated in the evolution of traits in cavefish that can be applied to other evolutionary genetic models.

## Introduction

Determining genotype-phenotype relationships is a central goal of biology, and critical for understanding how and why traits evolve. While significant advances have been made towards identifying the genetic variants contributing to natural variation in traits^1^, determining the genetic changes that underlie trait evolution remains challenging. In many species with derived traits, no closely related populations with the ancestral trait are extant, and the amount of genetic variation between phylogenetically distant populations makes it challenging to identify the genes and genetic variants associated with evolved traits. Moreover, many evolutionary models are not amenable to the use of the functional genetic tools available in other model organisms, and thus, functionally assessing candidate genes and genetic variants to demonstrate genotype-phenotype relationships is either time-intensive or not feasible. Thus, strategies are needed to both identify and functionally assess candidate genes for evolved traits in genetically tractable species.

Genetic mapping is a powerful method for identifying the genetic architecture contributing to evolved traits. In contrast to candidate gene-based approaches, genetic mapping approaches allow for screening for regions of the genome associated with natural variation in traits and are not reliant on prior knowledge of locus function. In closely related populations with distinct phenotypes, linkage mapping using methods such as quantitative trait loci (QTL) analysis has been widely used to identify loci associated with evolved traits (for example^2–4^). While this approach is a powerful method for identifying loci associated with complex traits, barriers to identifying causative genes and mutations underlying QTL remain. Quantitative trait loci frequently contain dozens to hundreds of genes, making it difficult to identify which gene or genes are causative. Moreover, the lack of tools for genetic perturbation in many evolutionary and ecologically relevant systems makes it challenging to assess the function of candidate genes and genetic variants.

One exception to this is *Astyanax mexicanus*, a species of freshwater teleost fish that has emerged as a powerful model for evolutionary genetic studies. *A. mexicanus* consists of two morphs: surface-dwelling fish that inhabit rivers and streams and blind cave-dwelling fish found in caves in Northeastern Mexico. At least two separate lineages of surface fish have colonized caves and given rise to over 35 populations of cavefish^5,6^. Mexican cave tetras have evolved numerous adaptations to living in the cave environment, including behavioral, morphological and physiological changes^7,8^. As cave and surface *A. mexicanus* are interfertile and live and breed in the laboratory, QTL analysis has been successfully performed in this species to identify the genetic architecture underlying and loci associated with cave-evolved traits^9–23^. Sequencing of multiple *A. mexicanus* genomes has allowed for the identification of numerous candidate genes underlying these QTL^24–26^. Additionally, the successful application of functional genetic tools in this species, including transgenesis and CRISPR-Cas9, now permits functional assessment of candidate genes^25,27–29^. Application of these approaches has successfully identified several genes contributing to trait evolution (for example^27–29^), however, as in other species, the genes and genetic variants underlying the vast majority of evolved traits in *A. mexicanus* cavefish remain unknown. Thus, efficient methods to identify and functionally test candidate genes are needed to harness the strengths of this system.

Here, we establish methods for identification and rapid functional assessment of evolved traits in *A. mexicanus* cavefish. We focus on eye degeneration, a trait that is a hallmark of cave species and which has been studied extensively in *A. mexicanus*^30–32^. We use a combination of datasets to identify strong candidate genes for eye degeneration in cavefish. Moreover, we develop a pipeline to rapidly assess candidate genes for roles in eye development and function using CRISPR-Cas9 gene editing. Together, these methods identify multiple genes that affect eye development in *A. mexicanus.* Further, this study provides a set of methods for overcoming two major barriers to establishing genotype-phenotype relationships in this species and others: identification of high-confidence candidate genes underlying QTL and rapid functional assessment of multiple candidate genes. These methods can now be applied to other cave-evolved traits in this species, as well as in other evolutionary model species.

## Results

### Using multi-method genomic evidence to identify 29 eye loss candidates

To identify genes associated with the evolution of eye degeneration in cavefish, we established a pipeline to both identify and functionally assess eye candidate genes in *A. mexicanus* (Fig 1A). To identify genes that contribute to eye degeneration and evolution in cavefish, we assessed genes that fell within eye quantitative trait loci (QTL) from previous studies^10,11,13–15^. We mapped markers flanking the QTL peaks for different eye-related traits – thickness of the different layers of the retina^13^, eye size^10,11,14,15^, and lens size^10^ – to the surface fish genome^25^ (AstyMex 2.0) to determine genes that fell under eye-related QTL. As these QTL contained hundreds of genes, we leveraged two other metrics. We determined if genes under QTL were differentially expressed between surface and cave fish in developing eyes by reanalyzing an existing RNA-sequencing dataset performed on eyes dissected from 54 hours post fertilization (hpf) surface fish and cavefish from the Pachón cave^33^. In addition, we determined if these genes showed evidence of selective sweeps in Pachón cavefish in our previously published analysis of sequenced wild-caught fish^34^. We identified 29 candidate genes for functional analysis that were under QTL, differentially expressed in eyes between cave and surface fish, and under selection in Pachón cavefish populations. Of the candidate genes, 23 were downregulated in eyes in cavefish relative to surface fish, while 6 of the genes were upregulated in cavefish compared to surface fish (Table 1, Figure 1A). Together, this establishes a method for combining metrics to identify high-evidence candidates for the evolution of cave-associated traits.

**Figure 1.**
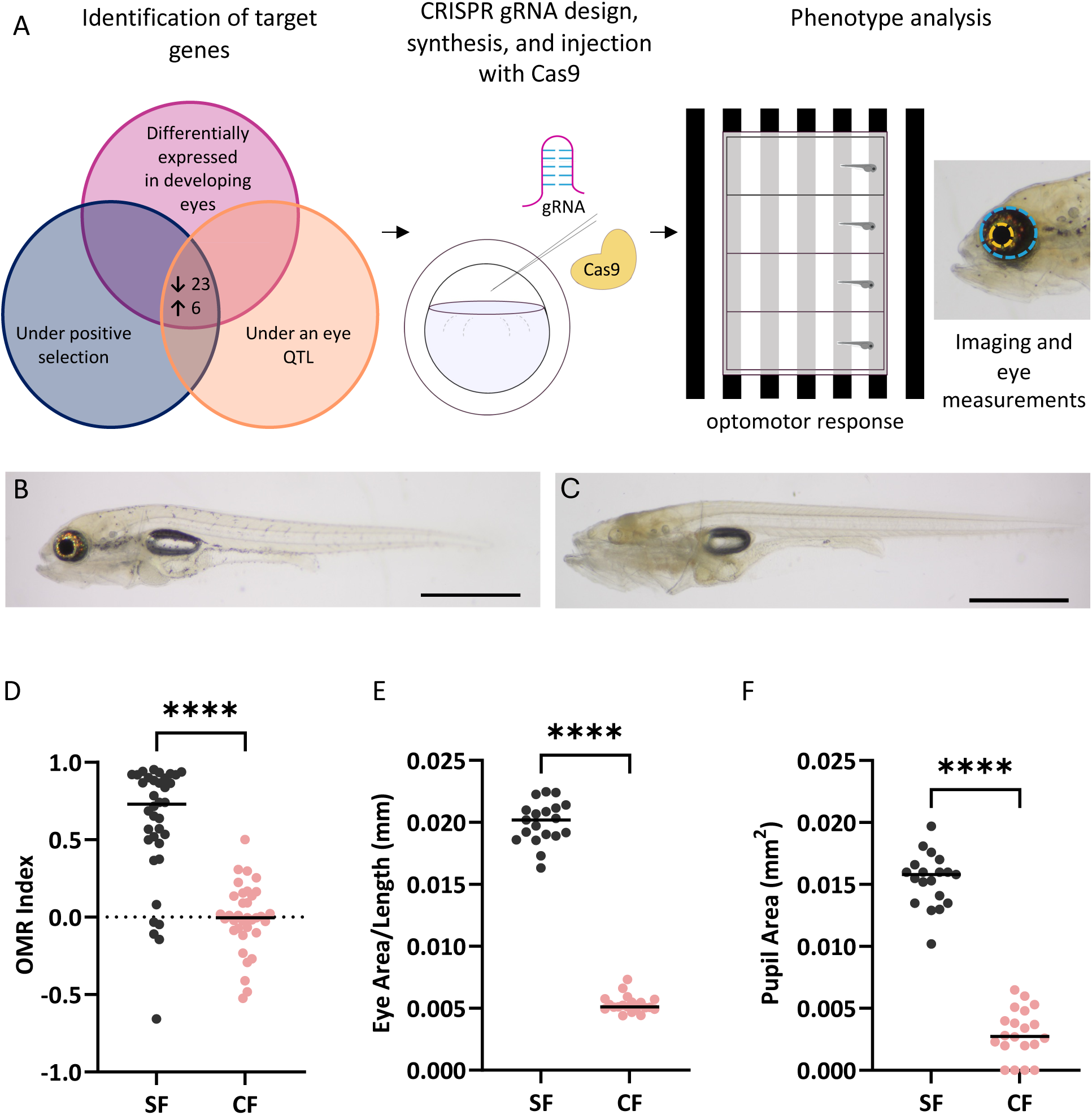
Pipeline for screening candidate eye genes. A. We identified candidate eye genes as those that met all three of the following criteria: 1) They were differentially expressed in Pachón cavefish eyes compared to surface fish eyes at 54 hpf in a previously published dataset^33^. 2) They fell under eye QTL derived from Pachón-surface hybrid crosses^10,11,13–15^. 3) They showed evidence of positive selection due to being present in selective sweeps in Pachón cavefish detected in population genetics datasets^34^. Only genes with an annotated gene name (29 genes) were assessed for further analysis. Following gene identification, we generated gRNAs to target each gene and injected individual gRNAs with Cas9 protein into single cell surface fish embryos. Injected, crispant fish and their wild-type siblings were assayed for optomotor response at 8 dpf, followed by eye and pupil size at 9 dpf. A subset of fish were analyzed for only eye/pupil size or OMR. Following phenotyping, fish were euthanized and processed for genotyping. B. Representative surface fish at 9 dpf. C. Representative Pachón cavefish at 9 dpf. D-F. Cavefish were compared to surface fish for all three phenotypes measured in the eye screen: D. Optomotor response, quantified as the OMR Index, in cavefish (n=35) compared to surface fish (n=36) (Mann–Whitney U test: p<0.0001). E. Standardized eye area in cavefish (n=20) compared to surface fish (n=19) (Unpaired t-test: p<0.0001). F. Lens area in cavefish (n=20) compared to surface fish (n=19) (Unpaired t-test: p<0.0001). Scale bars: 1 mm (B,C). \*\*\*\**p*<0.0001

**Table 1:**
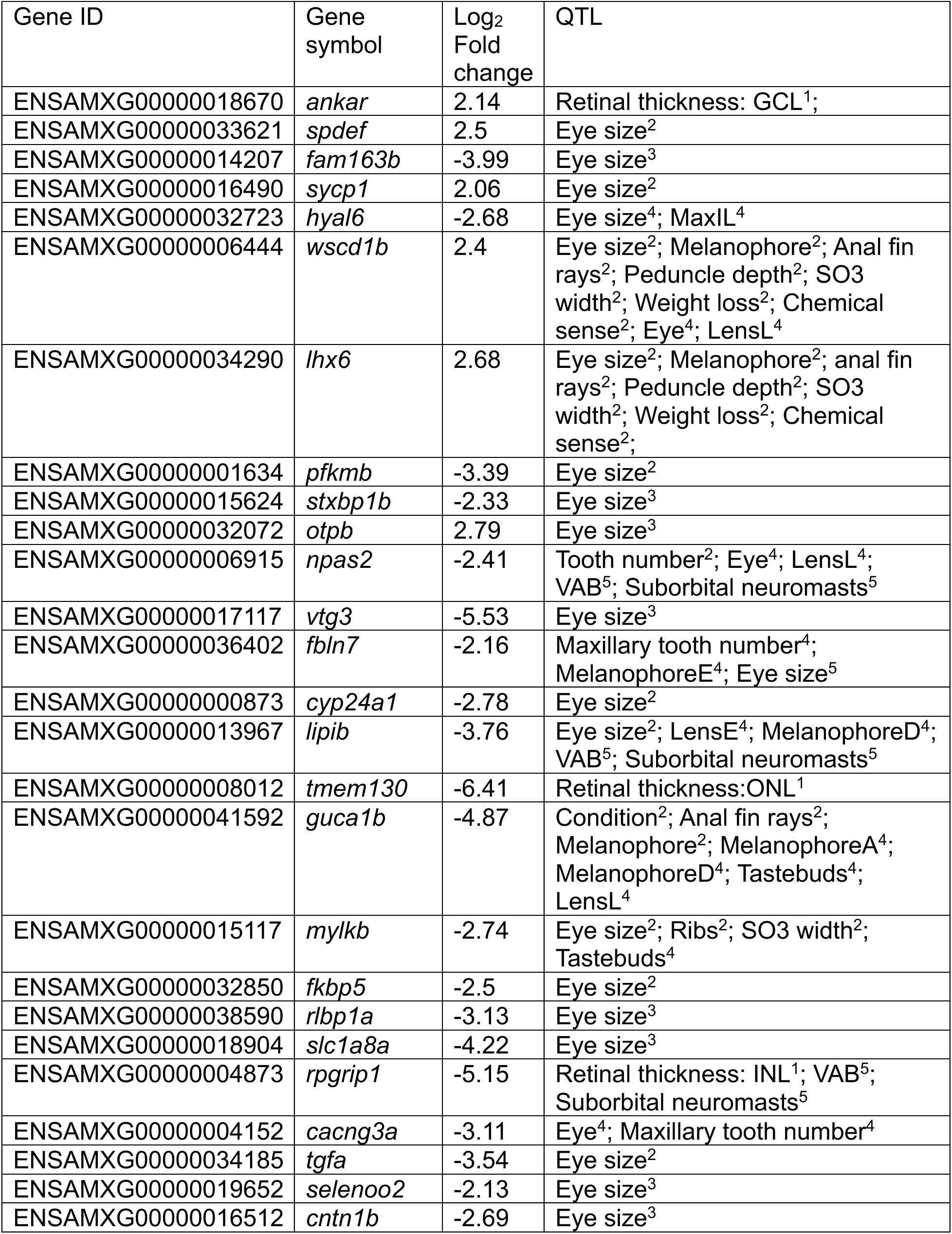
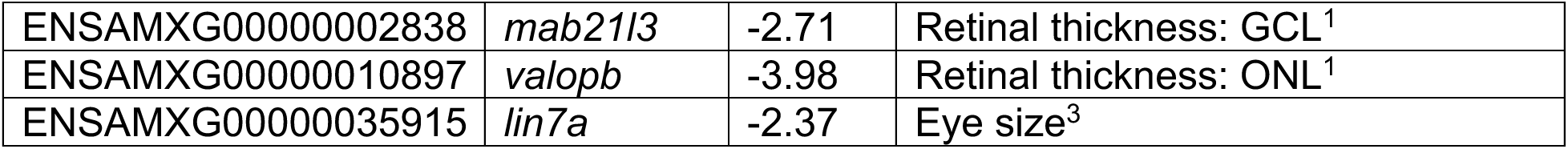
Candidate genes for evolution of eye loss. Genes were considered candidates if they met the following criteria: a) Differentially expressed in the eyes of Pachón cavefish and surface fish with a log_2_ fold change >2 b) Under an eye-related QTL from a surface-Pachón cavefish F2 hybrid cross. c) Evidence of hard or soft selective sweeps in Pachón cavefish. Gene IDs correspond to the surface fish 2.0 genome assembly. Fold changes are log_2_ transformed and indicate expression in cavefish developing eyes compared to surface fish developing eyes, such that a positive fold change indicates higher expression in cavefish compared to surface fish. QTL overlap was calculated by determining if the gene’s physical location occurs between the flanking markers of the QTL delineated in the original paper, using the surface fish genome (AstyMex 2.0). Traits with associated QTL are from the following: ^1^O’Quin et al. Quantitative genetic analysis of retinal degeneration in the blind cavefish Astyanax mexicanus. *PLoS One*. 2013. ^2^Protas et al. Multi-trait evolution in cave fish, Astyanax mexicanus*. Evol. Dev.* 2008. ^3^Kowalko et al. Convergence in feeding posture occurs through different genetic loci in independently evolved cave populations of Astyanax mexicanus. *Proc. Natl. Acad. Sci. U.S.* 2013. ^4^Protas et al. Regressive evolution in the Mexican cave tetra, Astyanax mexicanus. *Curr. Biol* 2007. ^5^Yoshizawa et al. Evolution of an adaptive behavior and its sensory receptors promotes eye regression in blind cavefish. *BMC Biol.* 2012.

### Analysis of eye candidate gene expression via single-nucleus RNA-sequencing reveals expression in eye-associated cell types

This method for identifying candidate genes did not rely on prior knowledge of gene function, suggesting that this method has the potential to capture candidate genes that were not previously known to play a role in eye development, a significant advantage over candidate gene approaches that require prior knowledge of gene function. To explore the potential role(s) of the candidate genes in development, we determined the transcriptionally defined cell types in which each of these genes is expressed in *A. mexicanus* surface and cave fish using single nucleus (sn)RNA-sequencing. We performed snRNA-seq on heads from surface fish and Pachón cavefish at 6 days post fertilization (dpf), a timepoint when larval fish begin to feed and perform many larval behaviors. We identified 35 clusters based on gene expression similarity (Fig 2A). Eye candidate gene expression was not restricted to a single cluster or group of clusters. While some genes, such as *fkbp prolyl isomerase 5* (*fkbp5)*, were expressed across many clusters, other genes showed more restricted expression. These included a set of genes, *phosphofructokinase, muscle b* (*pfkmb)*, *solute carrier family 1 member 8a* (*slc1a8a)*, *lipase member Ib* (*lipib)*, and *RPGR interacting protein 1* (*rpgrip1),* that were primarily expressed in cluster 8 which was enriched for genes that are expressed in photoreceptors (including *rcvrn2*, *gnat2*, *arr3a, gngt2b*). Other genes, including fibulin-7 *(fbln7*), *cytochrome P450 family 24 subfamily A (cyp24a1), neuronal PAS domain protein 2 (npas2*), *fkbp5*, *retinaldehyde binding protein 1a (rlbp1a*), *transforming growth factor α (tgfa*), and *synaptonemal complex protein 1 (sycp1),* were expressed in cluster 20, which was enriched for genes expressed in the retinal pigmented epithelium (including *rpe65a, rgrb, oca2*) (Fig 2B).

**Figure 2.**
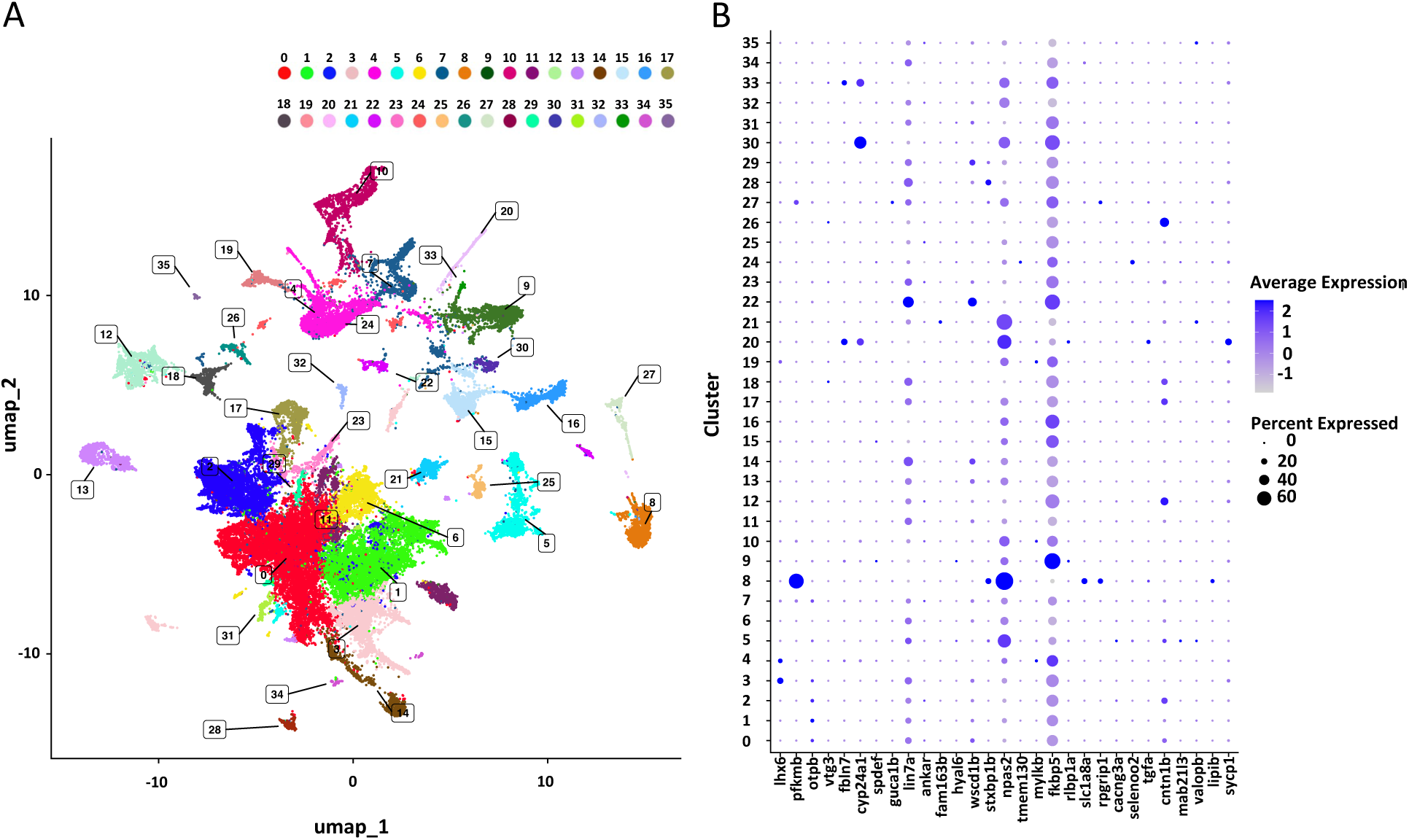
Analysis of eye candidate genes in single nuclei RNA-sequencing from larval heads. A. UMAP showing clusters obtained from single cell nucleus RNA-sequencing (snRNA-seq) of pools (n=15) of heads dissected from 6 dpf surface fish (n=2 pools) and Pachón cavefish (n=2 pools). The 36 clusters were manually annotated using enriched markers (see Supplemental Table 7). B. Dot plot to show the percent of cells expressing each eye candidate genes within each cluster (dot size) and the average expression of each eye candidate gene per cluster (dot color). Note that clustering and dot plots were generated for combined surface + cave nuclei. Genes *fbln7* = LOC103040604; *cyp24a1* = LOC103030576; *spdef* = LOC103037006; *fam163b* = LOC103027628 were relabeled with the gene names from the original surface fish 2.0 genome annotation in this figure to maintain consistency in gene naming across figures.

We determined whether each of the candidate genes was differentially expressed between surface and cave fish within each cluster. In many of clusters, none of the eye candidate genes showed differential gene expression when comparing surface to cave samples (Supplemental Table 1). In contrast, within the photoreceptor cluster, eight of the candidate genes were differentially expressed between cave and surface fish (Supplemental Figure 1, Supplemental Table 1). Together, these data suggest that the approach used here to determine candidate genes identified genes that are differentially expressed within multiple cell types in the eye. Moreover, many of these candidate genes are expressed within putative photoreceptors, suggesting they may alter photoreceptor development, maintenance, or function during eye degeneration in this species.

### Functional assessment of eye candidate genes using CRISPR-Cas9

To determine if each candidate gene plays a role in eye development, we developed a pipeline to functionally perturb eye candidate genes in *A. mexicanus* surface fish and assess eye-related phenotypes (Figure 1A). We used CRISPR-Cas9 to target each of the eye candidate genes and assess optomotor response (OMR), a behavioral readout of visual function during which fish follow moving lines ^35–37^, as well as two eye measurements, eye size and pupil size in injected fish, termed crispants (Figure 1A). These traits differentiate surface fish from Pachón cavefish during larval stages. Surface *A. mexicanus* have a robust optomotor response, indicated by an optomotor index around 1, whereas the optomotor response for cavefish is significantly reduced relative to surface fish, consistent with a lack of vision (Figure 1D). Both eye area and pupil area are significantly reduced in cavefish compared to surface fish (Figure 1B, C, E and F).

To verify that the crispant screening pipeline is sufficient for identifying differences in eye-related phenotypes in crispant fish, we tested our pipeline using fish in which *retinal homeobox 3 (rx3),* a gene that is required for eye development in a number of species^38–44^, including *A. mexicanus* ^29^, was targeted. The *rx3* crispant fish showed a range in eye phenotypes that distinguished them from wild-type fish, ranging from reduction in eye size to the complete loss of eyes and pupils (Figure 3A). Relative to wild-type, uninjected siblings, the average eye and pupil areas of the *rx3* crispant fish are significantly reduced (Figure 3B and C). Moreover, the *rx3* crispant fish have a significantly reduced OMR index compared to wild-type fish, with the median OMR index of *rx3* crispant fish around 0, suggesting that unlike their wild-type siblings, the majority of the *rx3* crispant fish do not respond to visual stimuli (Figure 3D). The results of these assays are consistent with the known function of *rx3* in *A. mexicanus*^29^, and establish that the crispant screening pipeline is a valid method to functionally interrogate eye candidate genes in this species.

**Figure 3.**
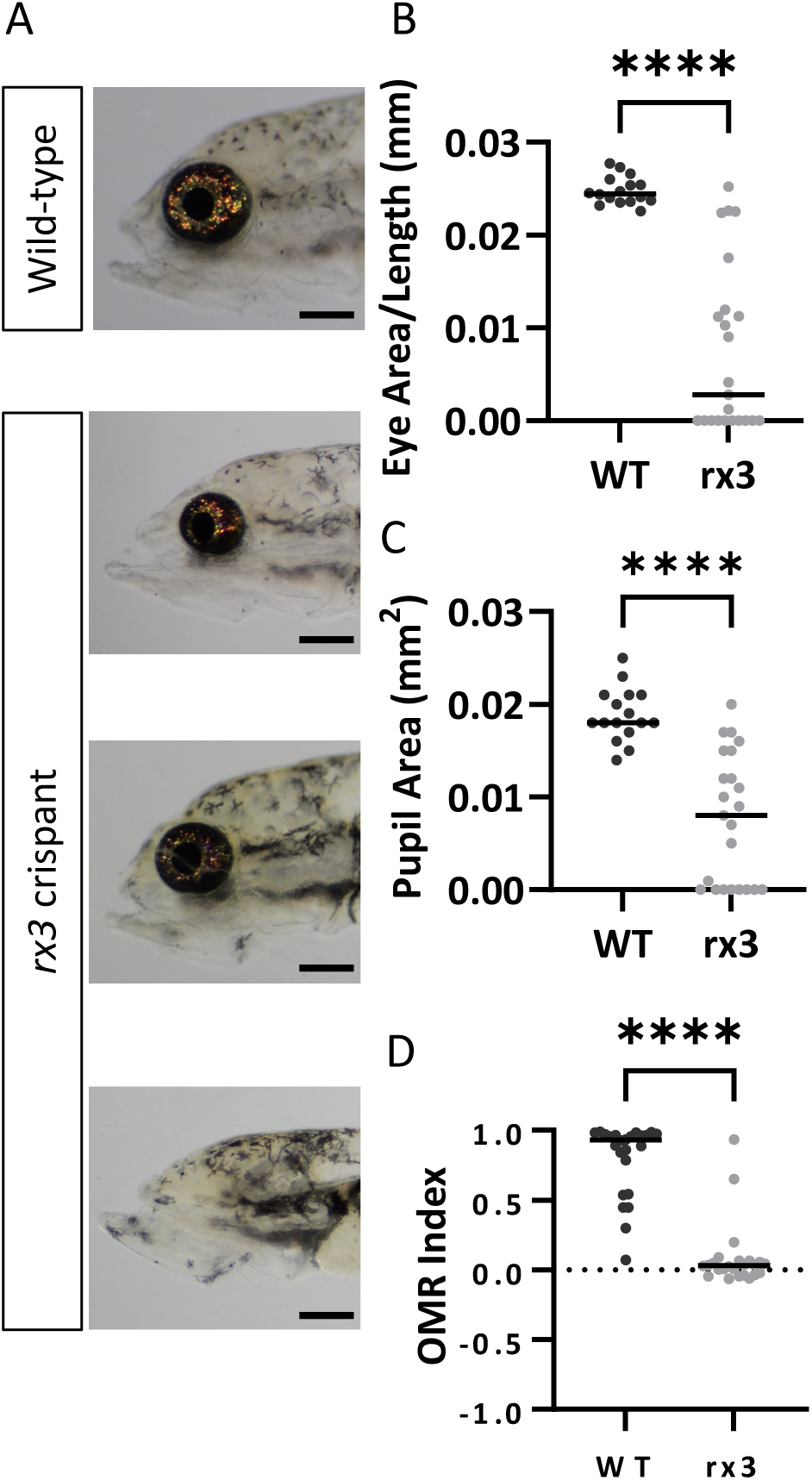
Phenotyping of *rx3* crispants establishes effectiveness of phenotyping pipeline. A. Images of wild-type sibling and *rx3* crispant surface fish at 9 dpf. B. Standardized left eye area in *rx3* crispant fish (n=23) compared to wild-type surface fish uninjected siblings (n=16) (Mann–Whitney U test: p<0.0001). C. Left pupil area in *rx3* crispant fish (n=23) compared to wild-type uninjected surface fish siblings (n= 16) (Mann–Whitney U test: p<0.0001). D. Optomotor response in *rx3* crispant fish (n=24) compared to wild-type uninjected surface fish siblings (n=24) (Mann–Whitney U test: p<0.0001). Scale bars: 0.5 mm (A). ****P<0.0001

### Crispant screening identifies multiple genes that contribute to variation in eye size

We used CRISPR-Cas9 to generate crispant surface fish with mutations in the eye candidate genes, successfully targeting 28 of the 29 candidate genes. As high rates of mutagenesis in crispant fish are essential to screen phenotypes in injected fish, which are mosaic for CRISPR-Cas9-induced disruptions targeted genes, we both verified mutagenesis qualitatively through PCR followed by gel electrophoresis, and quantified the per individual efficiency of each gRNA by performing amplicon sequencing targeting the gRNA target region within each gene from a subset of our 9 dpf crispant fish. The efficiency of mutagenesis was high for the majority of the gRNAs, with 18 out of 28 of the gRNAs producing an average mutagenesis of greater than 50%, and 11 of these gRNAs resulting in mutagenesis at an average of 80% or higher (Table 2, Supplemental Table 5). Moreover, efficiency was largely consistent across injected individuals targeted with the same gRNA (Supplemental Table 5). Together, these results support the use of this method to efficiently mutate candidate genes for quantitative phenotypic analysis.

**Table 2.**
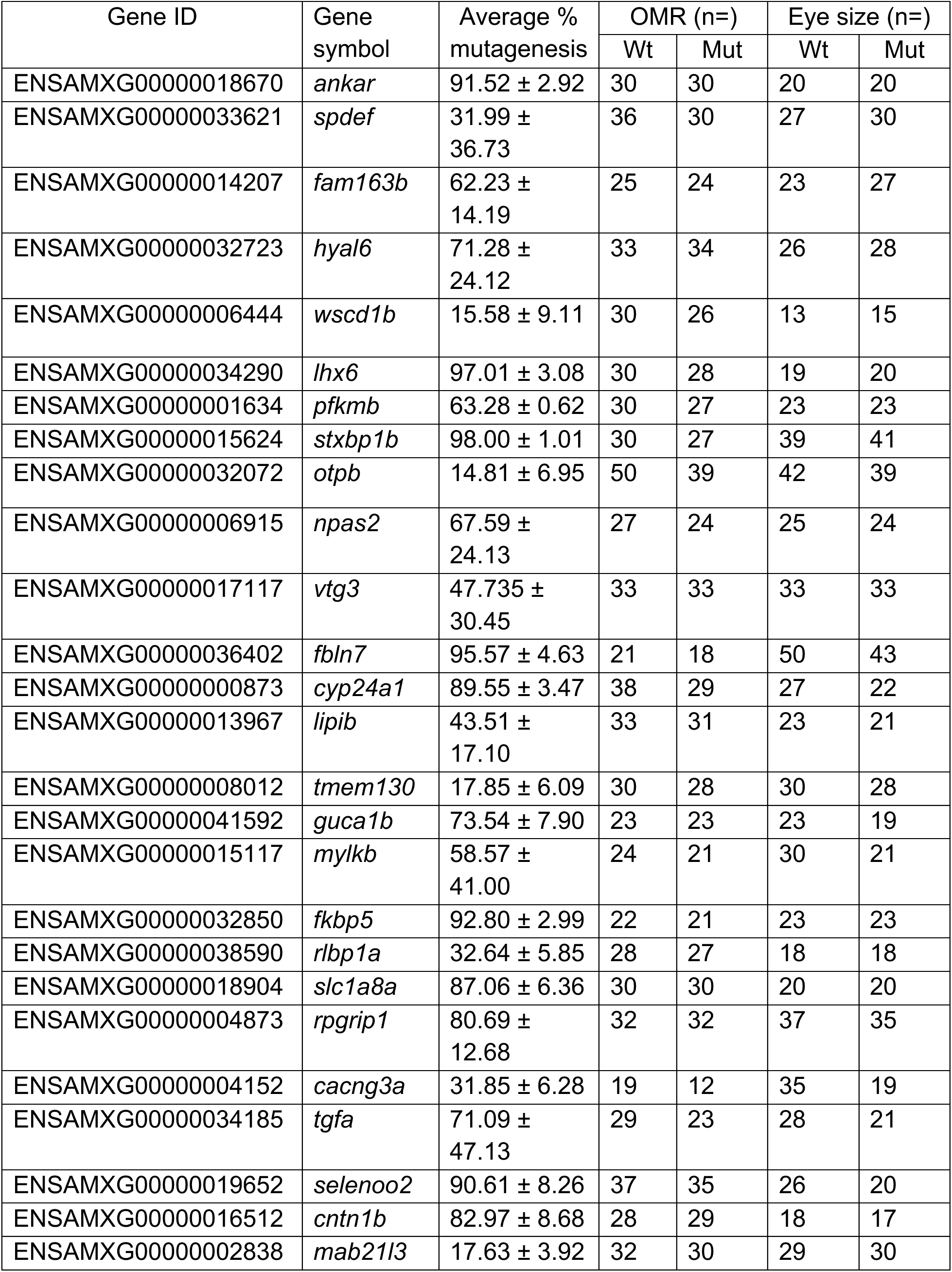
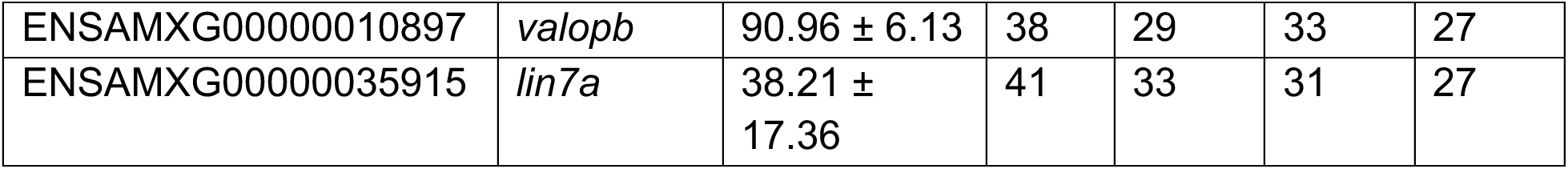
Eye screen statistics. Gene ID, followed by the gene symbol as found in the Ensembl version 2.0 surface fish genome. The average and standard deviation of percent mutagenesis per embryo, as calculated through percentage of modified alleles in Crispresso2^98^ following amplicon sequencing for crispants for each gene. Four larval fish were sequenced per gene. The number of wild-type and crispant fish assayed per gene in the primary screen are listed for each phenotype.

Eye size, pupil size and optomotor response were quantified and compared between crispant fish and their wild-type, uninjected siblings. Nine groups of crispant fish had statistically significant differences in left eye area compared to their wild-type counterparts (Figure 4A). Targeting the genes *orthopedia homeobox b* (*otpb), fbln7*, and *vitellogenin 3* (*vtg3)* resulted in a significantly reduced eye area in crispant fish compared to wild-type siblings. For the remaining eye candidate genes which impacted eye size, *lin7-homolog A (lin7a), SAM pointed domain-containing Ets transcription factor* (*spdef), cyp24a1, guanylate cyclase activator 1B (guca1b), LIM homeobox 6 (lhx6)*, and *pfkmb,* crispant fish had a increased eye area compared to wild-type surface fish siblings (Figure 4A, Supplemental Tables 3 and 4). In addition to eye size, left pupil size of *pfkmb* crispants was significantly increased compared to wild-type siblings (Figure 4B, Supplemental Tables 3 and 4). Results are largely similar for right eye and pupil areas (Supplemental Figure 2). For every gene, OMR indices were similar when comparing crispant fish to their wild-type siblings (Figure 4C, Supplemental Tables 3 and 4), suggesting crispant fish for all of these genes can still interpret visual cues. Overall, these data demonstrate that multiple eye candidate genes identified through our combinatorial approach regulate eye development. As all candidate genes fall under eye QTL, these data suggest that one or more of the candidate genes may play a role in the evolution of eye degeneration in cave *A. mexicanus*.

**Figure 4.**
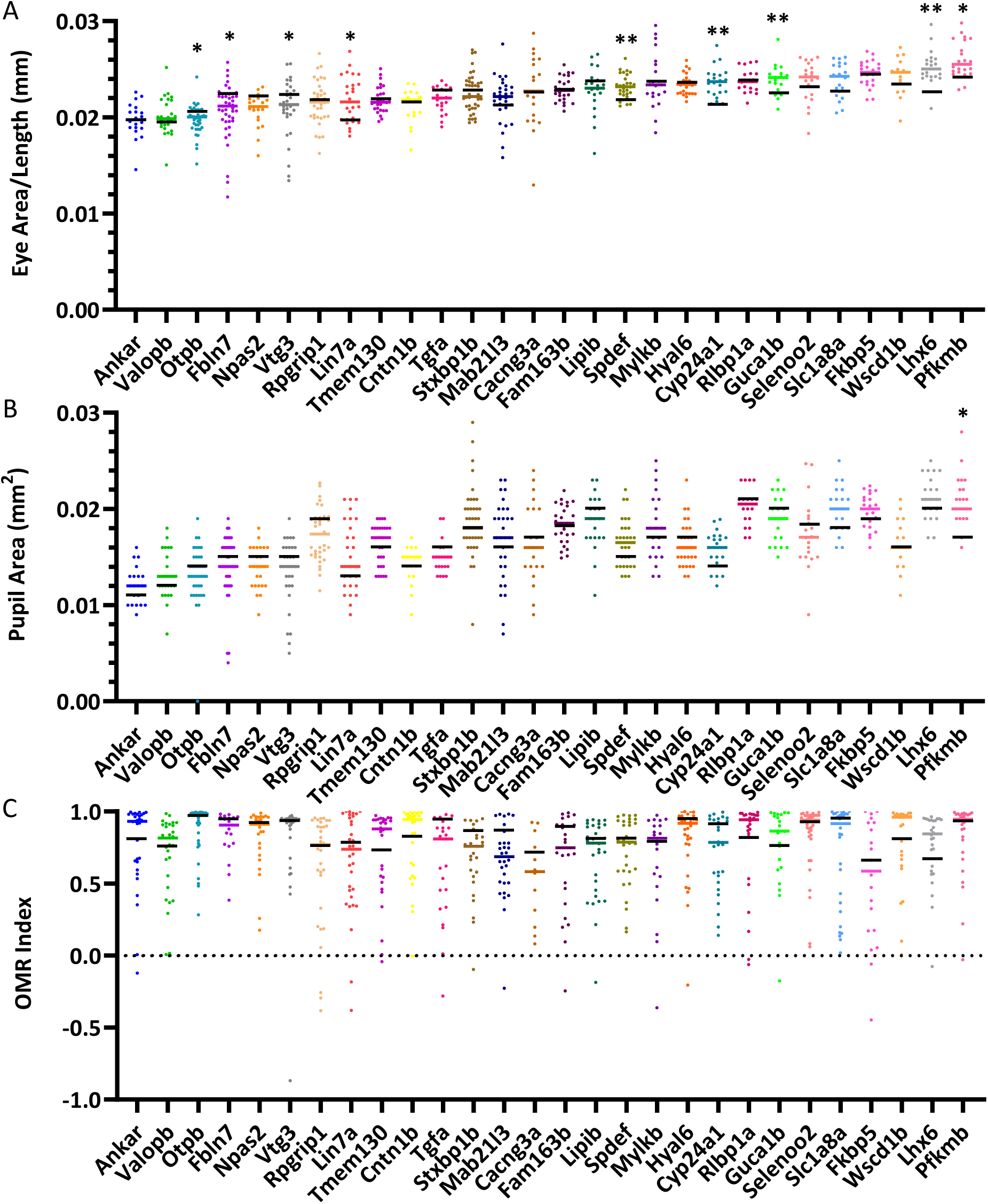
Crispant screening of eye candidate genes reveals regulators of eye and pupil size. A. Comparison of standardized left eye area between crispants of 28 genes and their wild-type uninjected surface fish siblings. Each colored data point represents eye size for 1 crispant fish, while the colored bar is the median eye size for crispant fish. Black bars are the medians of eye area for wild-type fish collected from the same clutch as injected crispant fish. (See supplemental tables 3&4 for measurements and statistics). B. Left pupil area in crispant fish compared to wild-type surface fish siblings. Each colored point represents pupil area in a single crispant fish and the colored bar is the median of the crispant pupil area. The black bar represents the median pupil area of the wild-type siblings. (See supplemental tables 3&4 for measurements and statistics) C. Optomotor response (OMR) index of crispant fish compared to wild-type siblings. All colored data points indicate OMR for 1 crispant fish, while the colored lines indicate the median OMR index of those crispant fish. The black bar is the median OMR index of wild-type siblings. (See supplemental tables 3&4 for measurements and statistics). Statistical significance was calculated following a correction for FDR. **Q<0.01, *Q<0.05

To confirm the findings in our screen, we targeted any gene which resulted in a phenotype in crispant fish with a second gRNA. The gRNAs used in this secondary screen targeted a different region in the gene. To account for potential effects of microinjections on eye size, crispant fish were compared to control fish which were injected with a scrambled gRNA which did not target any region in the *A. mexicanus* genome. Second gRNA injected fish and their scramble gRNA injected controls were imaged at 9 dpf to assess eye area. For two genes, *otpb* and *guca1b*, eye size of crispant fish did not follow the pattern observed in the primary screen, as medians for both scramble gRNA control fish and the second gRNA crispant fish were nearly indistinguishable (Supplemental Figure 3A and B). In contrast, crispant fish in which seven of the nine genes were targeted had eyes that trended in the same direction relative to control siblings as were observed in the primary screen for left and/or right eye: *fbln7*, *vtg3*, *spdef*, *lhx6, cyp24a1*, *pfkmb* and *lin7a* (Supplemental Figure 3A and B). The similar phenotypic results obtained by injecting two independent gRNAs further validate using crispant analysis as a method for phenotypic analysis of eye size. Together with results from the primary screen, these results suggest that we identified key candidate genes for eye degeneration in cavefish.

### Analysis of eye candidate genes in surface and cavefish suggests few coding variants between fish populations

One of the core challenges in evolutionary genetics is determining whether alleles underlying trait evolution exert their effects via coding sequences or regulatory elements. To determine if these genes harbor genetic variants that result in amino acid changes in cavefish, we assessed the sequences of the nine eye candidate genes which resulted in changes in eye size when mutated in surface fish. For each gene, we compared the predicted protein sequence between surface fish and Pachón cavefish using the published surface and cave fish genomes and assessed whether any variants that resulted in protein coding changes were present at high frequency in wild-caught Pachón cavefish and absent in two populations of wild-caught surface fish. This analysis included 10 individuals from Pachón cavefish populations and 10 individuals each from two surface populations: Río Choy and Rascón. We then compared the *A. mexicanus* surface and cave fish alleles with protein sequences derived from other fish species to investigate if any high frequency genetic variants were at conserved residues. Four of the eye candidate genes (*otpb*, *spdef*, *guca1b*, *lin7a*) did not contain any protein coding changes when surface and Pachón cavefish sequences were compared. In another four candidate genes (*vtg3*, *fbln7*, *pfkmb*, *lhx6*), we identified genetic variants that resulted in protein coding differences between surface and Pachón cavefish, however, in all of these genes the genetic variants were not exclusive to Pachón cavefish and were present at high frequency in one of the surface populations, or the Pachón cavefish alleles led to changes at residues that were not conserved across species (Supplemental Table 6). For a single gene, *cyp24a1*, we identified a single base pair change that led to a P->T amino acid change at a residue at which the surface allele is conserved across all fish species we assessed (Supplemental Table 6). These data suggest that protein coding changes within these genes are unlikely to contribute to the cavefish eye degeneration phenotype. Further, they support the role of regulatory changes, rather than protein coding changes, as being primarily responsible for phenotypic differences in *A. mexicanus* eye development.

### Mutation of the gene fbln7 impacts early stages of eye development

We further investigated the effects of perturbing one gene, *fbln7,* mutations in which resulted in reduced eye size in crispant fish during the primary screen (Figure 4A). The protein encoded by *fbln7* is known to play a role in angiogenesis^45^ and can inhibit neovascularization in the rat corneal angiogenesis model^46^. These data, along with the previously published work which shows defective optic vasculature at multiple time points during early cavefish eye development^28^, suggested that mutagenesis of *fbln7* may affect eye development at additional developmental time points. To determine if *fbln7* crispants affect eye size at early stages of eye formation, we targeted *fbln7* in surface fish with two gRNAs, and assessed eye development in crispant and wild-type fish at 2.5 dpf, and early stage of eye degeneration^30^. The *fbln7* crispant fish had significantly reduced eye areas compared to wild-type siblings at 2.5 dpf (Figure 5A and B). Pupil area was also reduced in *fbln7* crispants compared to wild-type, uninjected siblings at 2.5 dpf (Figure 5A and C). These data suggest that changes to early eye development in *fbln7* crispants may lead to the eye size reductions observed in larval stages in our screen. By 28 dpf, however, there are no quantifiable differences in eye size and pupil size between wild-type and *fbln7* crispant surface fish (Figure 5D-F). Together, these data suggest a role for the eye candidate gene *fbln7* in early developmental and/or early larval stages of eye development.

**Figure 5.**
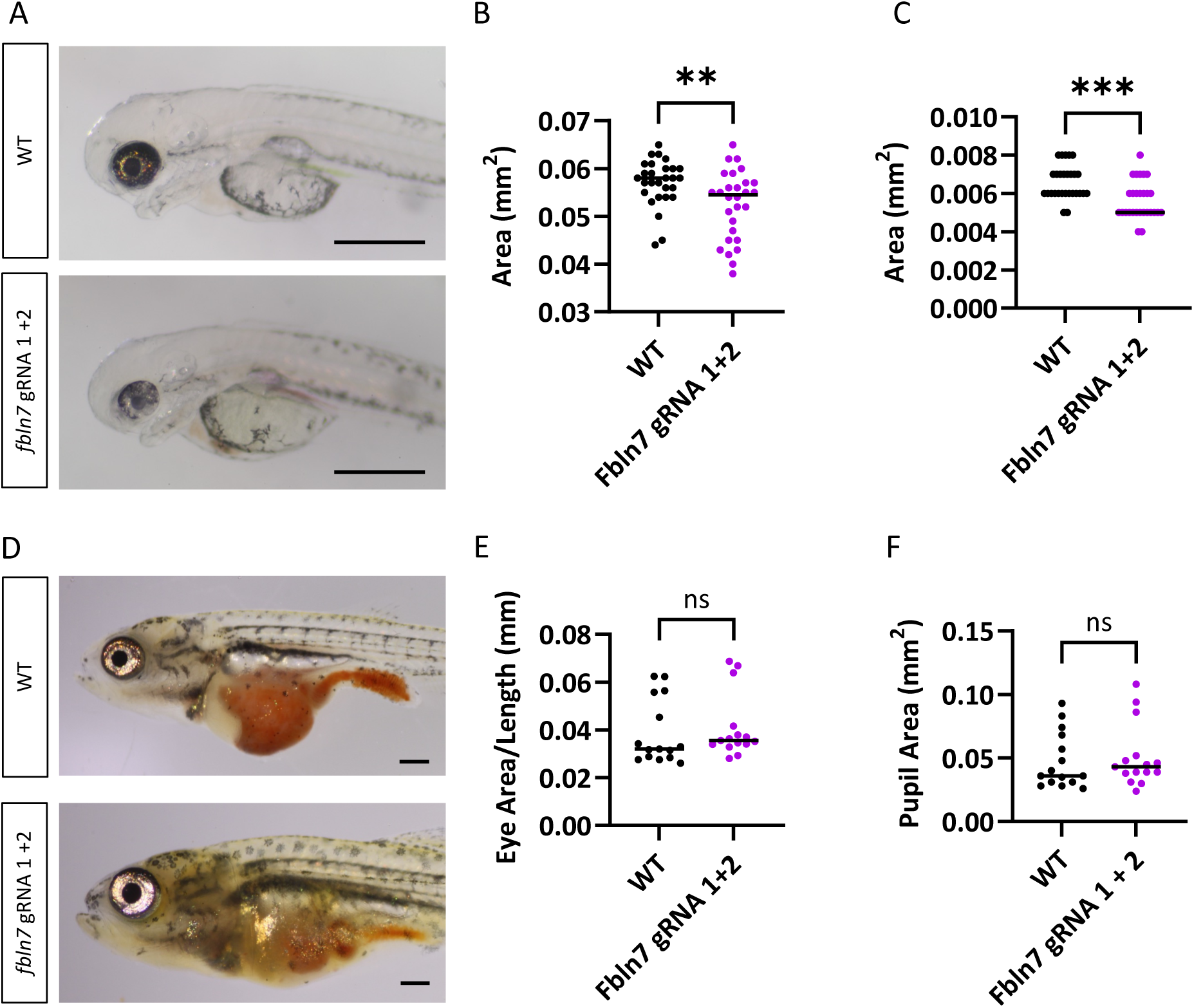
Eye area is altered during early stages of development in *fbln7* crispant fish. A. Images of 2.5 dpf wild-type surface fish (top) and *fbln7* gRNA 1 + 2 crispant fish (bottom). Scale bar = 0.5mm. B. Left eye area of *fbln7* gRNA 1+2 injected crispant fish (n=28) compared to wild-type siblings (n=28) at 2.5 dpf (Mann-Whitney U test: p<0.01). C. Left pupil area of *fbln7* gRNA 1+2 injected crispant fish (n=28) compared to wild-type siblings (n=28) at 2.5 dpf (Mann-Whitney U test: p<0.001) D. Images of 28 dpf wild-type surface fish (top) and *fbln7* gRNA 1+2 injected crispant fish (bottom). E. Standardized left eye area in 28 dpf wild-type fish (n=15) compared to *fbln7* gRNA 1+2 injected crispant fish (n=15) (Mann-Whitney U test: p=0.137). F. Left pupil area in 28 dpf wild-type (n=15) and *fbln7* gRNA 1+2 injected crispant fish (n=15) (Mann-Whitney U test: p=0.4923). B,C,E,F: Each data point represents the measurement for a single fish, and the colored line is the median.***=P<0.001 **=P<0.01

## Discussion

### Genetic underpinnings of eye loss in cavefish

Most cave-dwelling animals show reductions or loss of eyes^47^. In *A. mexicanus* cavefish, the processes that lead to eye development and ultimately degeneration have been studied extensively^31,48–54^. While a number of studies have investigated the genetic underpinnings of eye loss through genetic mapping studies^10,11,13–16,25^, the causative genes and genetic changes contributing to eye degeneration remain largely elusive. Exceptions to this are the *cystathionine β-synthase* (*cbsa)* gene, in which regulatory changes have been identified and likely contribute to eye degeneration in Pachón cavefish^28^ and the *rx3* gene, which is reduced in expression in multiple *A. mexicanus* cavefish populations in the early eye field^25,54,55^ and likely contributes to the evolution of reduced eye size via unidentified regulatory variants in Pachón cavefish^29^. Here, we identified an additional nine genes which fall under eye QTL, are differentially expressed in cave and surface fish whole eyes and/or populations of cells, exhibit signatures of selection in cavefish, and alter eye size when mutated with CRISPR-Cas9 in surface fish.

Recent work has investigated whether the many of the genes that contribute to eye size in *A. mexicanus* do so through regulatory, rather than coding changes. A lack of protein coding variants in *cbsa* and *rx3* and evidence of cis-regulatory differences between cave and surface fish at each of these loci is consistent with cis-regulatory variation affecting eye size in cavefish through both of these genes^28,29,55^. Moreover, assessment of a list of known eye-related genes across cavefishes revealed that few vision associated genes have putative non-functional alleles in *A. mexicanus* cavefish^56^. In contrast, ten eye-related genes contained premature termination codon at high frequences in *A. mexicanus* cavefish populations but not in surface fish^57^. For one of these genes, *phosphodiesterase 6C* (*pde6c),* functional perturbation in surface fish resulted in loss of vision^57^. Thus, a combination of coding and regulatory variants likely contribute to evolution of eye loss in *A. mexicanus* cave fish. Here, we find that of the nine candidate genes which result in changes in eye size when functionally manipulated, only half contained coding variants in Pachón cavefish relative to surface fish, and the majority of the identified coding variants are at non-conserved residues when these genes are assessed across species. The exception to this is *cyp24a1*, in which a variant in Pachón cavefish causes an amino acid change relative to surface fish at a residue which is conserved across species. Together, these data suggest that if these eye candidate genes have genetic variation that is contributing to the Pachón cavefish eye phenotype, the majority of the causative alleles are likely regulatory in nature.

Of the nine genes that altered eye size in our primary screen, we investigated *fbln7* across multiple developmental stages. At 6 dpf, *fbln7* expression was observed in a subset of cells in only a few clusters. One of these is cluster 20, the cells of which are hypothesized to be a part of the retinal pigmented epithelium (RPE). The RPE is a pigmented cell layer outside photoreceptor layer of the retina which plays a number of essential roles in eye development and vision^58^ and has been shown to play a role in eye degeneration in *A. mexicanus* cavefish^59^. Within cluster 20, expression of *fbln7* is reduced in cavefish compared to surface fish, although this reduction does not reach statistical significance (p=0.09). These findings, combined with the reduction of eye size in *fbln7* crispant fish at multiple stages of eye development, raises the possibility that *fbln7* plays a role in eye degeneration in cavefish via a function in the RPE.

### CRISPR screening is a powerful method for identifying the genetic basis of trait evolution

While the causative genetic variants contributing to trait evolution have been under investigation for decades and progress has been made towards identifying genetic variants contributing to trait evolution in a variety of species^1^, identifying and functionally assessing genes associated with trait evolution remains challenging. A central challenge is devising methods to rapidly assess many candidate genes. CRISPR-Cas9 can now be applied to a variety of model systems (for example^60–62)^, providing a platform for functional testing of candidate genes.

In zebrafish, CRISPR screening has helped determine a number of key genes in a variety of processes, including regeneration in the retinal pigment epithelium^63^ neurobehavior^64^, and heart regeneration^65^. In non-model systems, work has been done to establish CRISPR-based mutagenesis in a number of evolutionarily interesting species, including the beetle *Tribolium castaneum*^66,67^, stickleback fish *Gasterosteous aculeatus*^68^, and the amphipod crustacean, *Parhyale hawaiensis*^69^. However, these tools have largely been used to target individual candidate genes in these species. Similarly, in *A. mexicanus,* CRISPR-Cas9 was first applied to investigate the role of an individual candidate gene, *oca2*, in the evolution of albinism^27^. While CRISPR has now been used to mutate candidate genes for both morphology^29^ and behavior^70–72^ in *A. mexicanus*, these past studies have used CRISPR to investigate one or a small number of genes hypothesized to be involved in these traits. Here, we show that CRISPR can be utilized in *A. mexicanus* to perform screening of dozens of genes in a high-throughput manner. This approach is critical, as it alleviates the need to prioritize genes based on known gene function. Indeed, a subset of the eye candidate genes we screened in this study had no known role, to our knowledge, in eye development or maintenance. These included genes, such as *otpb* and *vtg3*, which ultimately resulted eye size phenotypes when perturbed in *A. mexicanus* surface fish. Thus, this approach allows us to rapidly assess a large number of genes using CRISPR-Cas9, permitting the testing of candidate genes which may not be targeted using other approaches.

While we only assessed eye and vision-related phenotypes here, a number of other morphological, physiological, and behavioral traits differ between cave and surface *A. mexicanus*^5,16,17,73–84^. Genetic mapping has been utilized to investigate the genetic underpinnings of many of these traits, including reductions and loss of pigmentation, changes in metabolism, and alterations to feeding behavior including feeding posture and vibration attraction behavior^9–11,14–17,25,26,85,86^. The methods used for both narrowing lists of candidate genes and rapid functional assessment of candidate genes described here can be applied to investigate other cave-evolved traits in this species. In addition, the efficiency of mutagenesis found here suggests that this method will be feasible to assess traits that manifest at later stages of development. Finally, in a number of evolutionary model systems, genetic mapping is used to investigate the genetic underpinnings of trait evolution and gene editing is feasible (for example the African turquoise killifish, *Nothobranchius furzeri*^87,88^, sickleback fish, Gasterosteous aculeatus^68^, and cichlids^89^). Thus, these methods have broad applicability to other evolutionary model organisms.

## Materials and Methods

### Selection of candidate genes

To determine if genes fell under eye QTL, markers flanking previous published QTL for the eye-related traits retinal thickness^13^, eye size^10,11,14,15^, and lens size^10^ were mapped to the *Astyanax mexicanus* surface fish genome (AstyMex 2.0)^25^. RNA-sequencing data previously performed on developing surface and cave fish eyes at 54 hpf^33^ was mapped to the surface fish genome (AstyMex 2.0)^25^ and evaluated for differential expression. Genes which fell under QTL and were differentially expressed were evaluated for signatures of selective sweeps using diploS/HIC^90^ on sequencing from wild-caught individuals^5,34^, as done previously^34^. Genes which fell within QTL, showed differential expression greater than log 2 fold in surface fish compared to cavefish eyes, and which showed signatures of positive selection, classified as hard or soft sweeps using diploS/HIC^90^ in the Pachón cave population, were considered candidates (Fig 1A, Table 1). Any genes which were unannotated or which were not annotated with gene names were excluded.

### Fish husbandry and breeding

All *A. mexicanus* fish used in this study were generated from lab-bred stocks that originated in Mexico and were maintained at Lehigh University fish facilities. All fish husbandry and handling were performed in accordance with Lehigh University Institutional Animal Care and Use Committees (IACUC) approved protocols. All *A. mexicanus* fish were kept under a 14:10 light/dark cycle. Larvae were raised in glass bowls at 25°C in densities of approximately 100 fish per bowl until 6 days post fertilization (dpf). At 6 dpf, fish were transferred to 2-liter tanks at densities of 30 fish per tank. Juvenile and adult fish were housed at 23 +/- 1°C. Beginning on 6dpf, fish were fed brine shrimp. Adult fish that were utilized for breeding received bloodworms and pellets (Zeigler Zebrafish Diet SKU # AH271, Pentair). To induce spawning, bloodworms were fed with greater frequency, and the temperature of the tank was raised by ∼2°C via submersible heater.

### Generation of crispant fish

CRISPR gRNA target regions were chosen in each gene using CHOPCHOP^91^. The gRNA target sequences that were preferentially selected were predicted to have a high efficiency, few to zero off target sites, and two five prime G nucleotides (Supplemental Table 2). gRNAs were synthesized as previously described ^25,27,92^. Single cell surface fish embryos were injected with 1 nL of injection mix for a final concentration of 25 ng/mL gRNA and 100 ng/mL of Cas9 protein (PNA Bio, CP01-50). Sibling, uninjected single cell embryos were collected concurrently to serve as controls. For the majority of genes, fish from multiple clutches, injected on multiple nights, were combined. Of the 29 genes, we were able to successfully mutate 28 genes. An exception to this was the gene *sycp1*, which was not successfully mutated after 5 different gRNAs were injected.

A second gRNA was designed to target a different region of candidate genes which, when initially targeted, produced a phenotype in the assays. For this secondary screen, a gRNA using a scrambled sequence that does not target the *A. mexicanus* genome was generated^93^ (Supplemental Table 2) and injected with Cas9 protein as described above and these injected fish were used as controls.

### Optomotor response assays

At 8 dpf, crispant fish (fish injected with a gRNA targeting a gene of interest) and control uninjected siblings or wild-type surface and cave fish were screened for visual defects using an optomotor response (OMR) assay, as in^57^. Prior to the assay, the fish were fed and then placed in the assay room for at least one hour to acclimate. For each trial, fish were transferred to a 4-well rectangular plate (Nunc rectangular dishes, Thomas Scientific, item number 1228D90) filled 90% full of fish system water and placed on top of a Samsung Tab Active Pro tablet (model number SM-T540), with one fish per well. Fish were acclimated to a white background for 1 minute. Each trial began with 30 seconds of a white background, followed by a display of moving repeated black and white vertical lines that were 2 centimeters thick and moved at a speed of 1 centimeter per second. The moving lines switched direction every 30 seconds for 5 total switches and ended with 30 seconds of white background. Assays were recorded from above with a FLIR Grasshopper3 High Performance 329 USB 3.0 Monochrome Camera (Edmund Optics Cat. No. 88-607) with a 12 mm HP series lens, 330 1/1” (Edmund Optics, Cat. No. 86570) at 30 frames per second at a resolution of 800 x 1200 using the program Spinview from FLIR’s Spinnaker SDK.

Videos were analyzed to determine the distance that each fish moved in the direction of the moving lines as previously described^57^. Briefly, the distance traveled was measured by recording the x coordinate of the fish at the end of each line direction change using the program Fiji^94^. The x values before and after each line shift were subtracted from one another to produce the distance the fish swam relative to the direction of the lines. The total distance traveled in the direction of the lines was divided by the total possible distance (the arena length) to determine an OMR index score for each fish. The OMR index score ranges from 1 to −1, where −1 indicates that the fish traveled the total distance of the area in the direction opposite of the moving lines after each line direction change, while 1 indicates the fish traveled the entire distance of the arena in the direction of the moving lines after each line direction switch. Immediately following the assay, individual fish were placed in fish water in 12-well plates and then imaged and genotyped the following day. As groups of wild-type fish occasionally did not respond to the assay, for any OMR assay (a group of trials on the same day on the same clutch of fish) for which 80% of wild-type fish did not obtain an OMR index of 0.4 or higher, the entire assay was excluded from analysis. A subset of the fish assayed for OMR were imaged for eye size the following day (see below).

### Imaging

Eye phenotypes were assessed in wild-type and crispant surface fish via imaging at 9 dpf. Fish were euthanized in a solution of buffered tricaine methanesulfonate (MS-222). Fish were mounted in methylcellulose and images were taken with a Canon Rebel DSLR camera on a Zeiss Stemi 508 dissecting microscope. Images were analyzed by measuring eye area, pupil area, and body length (standard length) in Fiji^95^ (Fig 1A). In wild-type surface fish, eye area correlated with standard length. To account for the effect of body size on eye area, eye area was normalized by dividing eye area by standard length for comparisons. In wild-type fish, pupil area did not correlate with standard length, so pupil area not corrected for standard length. After imaging, all injected fish and a subset of wild-type controls were processed for DNA extraction and genotyping.

### Statistical analysis for phenotyping

Fish that had a small or uninflated swim bladder was excluded from analyses. Statistical analyses were conducted using GraphPad Prism version 10.6.1 (GraphPad Software, www.graphpad.com.). All data were evaluated for normality and equal variance. Unpaired student’s t-test was used when the data were normally distributed and a non-parametric Mann-Whitney U test was performed for data that were not normally distributed.

For data shown in Figure 3 and Supplemental Figure 2, crispant fish were compared to sibling, wild-type uninjected fish using a non-parametric Mann-Whitney U test. Following this comparison, a False Discovery Rate (FDR) analysis was performed to assess Type I errors. *P*-values were indicated as follows: *p < .05; **p < .01; ***p < .001, ****p < .0001.

### Genotyping

To identify mutagenic individuals, genotyping was performed as previously described^92,96,97^. Briefly, fish were placed into 100ul of 50 mM NaOH following euthanasia and incubated at 95°C for 30 minutes to extract DNA. 10ul of 1 M Tris-HCl pH 8 was added to each sample post incubation. Primers were designed to amplify the region around gRNA target sequences (Supplemental table 2). PCR was performed using the DNA extracted from every injected individual. Mutagenesis was assessed via gel electrophoresis as previously described^92,96,97^. When a gRNA did not produce mutagenesis that was visible by this method, a second gRNA was designed to the gene of interest. Mutagenesis was then confirmed and quantified using amplicon sequencing (see below).

### Assessing gRNA efficiency via amplicon sequencing

To quantify mutagenesis in injected fish, amplicon sequencing was performed to assess how efficient CRISPR-Cas9 mutagenesis was for each gRNA. PCRs were performed to amplify the regions surrounding the gRNA target site on four fish per gene, all of which had previously been confirmed for mutagenesis via PCR and gel electrophoresis. Following PCR using gene specific primers (Supplemental Table 2), a PCR cleanup was performed using a QIAquick PCR & Gel Cleanup Kit (Qiagen, 28506). Amplicon sequencing was performed (AmpSeq, Maryland). Briefly, libraries were prepared using the NEBNext Ultra II DNA Library Prep Kit for Illumina using the TruSeq Index Primer. Sequencing was performed using the Element Biosciences AVITI platform with the 2 x 150 cycle sequencing kit. Sequencing results were analyzed via CRISPResso2^98^ to determine the percentage of reads that were modified versus unmodified for each injected fish (Table 2, Supplemental Table 5).

### Assessment of coding variants between cave and surface fish

To determine if cavefish harbor coding variants in the candidate genes which produced eye phenotypes when targeted with CRISPR-Cas9, two methods were used. Protein sequences were derived from coding sequences of each gene as annotated in NCBI and Ensembl Beta in the surface fish genome (AstMex3) and the Pachón cavefish genome (AMEX_1.1). As the original gene list was derived from the Astyanax mexicanus 2.0 surface fish genome, BLAST of the surface fish Astyanax 2.0 coding sequence was used to ensure that all coding sequences assessed were for the equivalent gene targeted using CRISPR. Surface and Pachón cavefish protein sequences were aligned in AliView^99^ and inspected manually for differences.

In parallel, we determined if observed protein coding variants which distinguished Pachón cavefish from surface fish were at high frequency in surface versus cavefish populations using previously published sequencing data from wild-caught individuals^5^. Coding sequences for 10 Pachón cavefish and surface fish from Río Choy (n=10) and Rascón (n=10) were isolated through alignment to the *Astyanax mexicanus* 2.0 surface fish genome annotations from Ensembl, aligned in AliView^99^, and assessed manually. Protein coding changes that were present in at least 80% of Pachón cavefish individuals and which were not present in either surface fish population were considered high frequency derived alleles.

Any genes harboring variants that were predicted to cause protein coding changes and were high frequency in Pachón but were not present in surface fish populations were assessed to determine if the amino acid residue present in surface fish was conserved across species. We obtained predicted protein coding sequences for 14 Actinopterygian fish species (excluding *Astyanax mexicanus*) from the UCSC 100-way vertebrate multiz alignment to the human genome (hg38)^100,101^. We identified the human ortholog of each gene in zebrafish, using data from Ensembl BioMart and the Matched Annotation from NCBI and EMBL-EBI (Mane) Project^102^. We then obtained the coding sequences from the predicted protein alignments for the canonical Ensembl transcript (*cyp24a1* and *fbln7*) or from the predicted exon alignments for the NCBI RefSeq transcript (*pfkmb*) or the human ortholog (see transcript IDs in Supplementary Table 6). For *vtg3*, for which there was no human ortholog, we obtained sequences for the 7 Actinopterygian fish within the UCSC 8-way fish multiz predicted exon alignments, using the Ensembl transcript from Fugu^100,101^. We added the predicted coding sequences of these genes from the *Astyanax mexicanus* surface fish (AstMex 3) and Pachón cave fish (AMEX_1.1) reference genomes in NCBI, then translated all sequences to amino acids, and aligned the predicted amino acid sequences using Muscle^103^ in Seaview^104^ version 5.0.4. We then manually inspected the alignments at the position of variants of interest for conservation across other fish species.

### Tissue collection for single nuclei RNA sequencing

Six-day post fertilization surface fish and Pachón cavefish were euthanized in MS-222 and decapitated. Heads were flash frozen in pools of 15 in liquid nitrogen and maintained at −80C until processing at the University of Houston Sequencing and Gene Editing Core. Two pools of surface fish heads and two pools of Pachón cavefish heads were processed for single nuclei RNA sequencing.

### Single-nuclei RNA Library Preparation and Sequencing

Nuclei were isolated using a two-stage protocol that combines mechanical disruption with immunomagnetic purification at the University of Houston Sequencing Core. Single-nuclei suspensions were generated using Nuclei Extraction Buffer (Miltenyi Biotec) for rapid and gentle dissociation of all four samples. Samples were transferred into pre-cooled gentleMACS™ C Tubes (Miltenyi Biotec) containing ice-cold lysis buffer supplemented with RNase inhibitor (0.2 U/µL) to prevent RNA degradation. Mechanical dissociation was performed on a gentleMACS Dissociator (Miltenyi Biotec) using a cold program for five minutes. The resulting crude nuclear suspension was filtered through a 70µm MACS® SmartStrainer (Miltenyi Biotec) to remove aggregates and large debris, followed by washing in lysis buffer and centrifugation (300 × g, 5 min, 4 °C) to pellet nuclei. The pellet was resuspended in ice-cold nuclei separation buffer (PBS, pH 7.2, supplemented with MACS BSA Stock Solution at 1:250 and Nuclei Extraction Buffer at 1:7) and then filtered through a 30 µm SmartStrainer to ensure a homogeneous single-nucleus suspension.

Nuclei were magnetically labelled with Anti-Nucleus MicroBeads (Miltenyi Biotec) for enrichment and debris removal. One million nuclei were resuspended in 450 µL of nuclei separation buffer, incubated with 50 µL of MicroBeads for 15 min at 2–8°C, and diluted with 2 mL of buffer prior to separation. The labelled suspension was passed to an LS Column (Miltenyi Biotec) positioned within a MACS Separator (Miltenyi Biotec). Flow-through containing debris and unlabeled material was discarded, and the column was washed twice with 1 mL of separation buffer. Retained nuclei were then eluted from the column after removal from the magnetic field by flushing with 1 mL of separation buffer.

This nuclei suspension was diluted to reach 700-1200 nuclei/µL and used as an input for automated microfluidic single nuclei capture and barcoding using the 10X Genomics Chromium IX platform. snRNA-seq libraries were generated using Chromium NEXT-GEM Single Cell 5′-HT Gene Expression kit (10X Genomics) according to the manufacturer’s protocol at the University of Houston Sequencing Core. Single-nuclei gel beads in emulsion (GEMs) were generated, and single nuclei were uniquely barcoded. cDNA was recovered and selected for using DynaBead MyOne Silane Beads (Thermo Fisher Scientific) and SPRIselect beads (Beckman Coulter). This sequence library underwent quality control assessment using a High-sensitivity d1000 DNA screentape (Agilent) and then quantified with a Qubit Fluorometer (Thermo Fisher Scientific) and with a Kapa Library Quantification Kit (Kapa Biosystems) using the AriaMX instrument (Agilent). Sequencing was performed on an Illumina NovaSeq X+ system with the pair-end sequencing settings Read1 – 28 bp, i7 index – 10bp, i5 index – 10 bp and Read2 – 90 bp.

### Single-nuclei RNA Data processing and Analyses

The 10x Genomics Cell Ranger v8.0.1 using 10x Genomics Cloud Analysis^105^ was used for data demultiplexing, transcriptome alignment, and UMI counting. A custom *Astyanax mexicanus* reference genome was generated using Cell Ranger mkref with NCBI genome assembly GCF_023375975.1 and corresponding gene annotation file obtained from NCBI. The downstream analysis was performed in R software (v4.4.2), using the Seurat package^106^ (v5.3.1). First, the nuclei for each object (2 surface fish and 2 Pachón cavefish) were filtered based on unique feature counts, with the exclusion criteria of nuclei containing less than 300 and more than 5,000 unique features. The filtered nuclei were further refined by identifying and excluding doublets using the DoubletFinder package^107^. Nuclei from each Seurat object were separately normalized using SCTransform function and subjected to Principal Component Analysis using RunPCA function. FindNeighbors and FindClusters functions were used for clustering nuclei and visualized using RunUMAP function (Uniform Manifold Approximation and Projection). To eliminate the batch effects, the individually processed Seurat objects were integrated using the SCTIntegration workflow. The integrated object was used for cell type identification of nuclei at a cluster resolution of 0.2. Unbiasedly obtained clusters were identified by analyzing known function and expression of genes enriched in each cluster (Supplemental Table 7). The 29 genes of interest were investigated for their expression patterns using dot plots in R. Further, differential expression analysis was performed for the subset of 29 genes to determine if each gene was differentially expressed between surface and cavefish in each cluster. Note that four genes which were annotated with names in the original surface fish genome dataset were annotated differently in the AstyMex3.0 dataset. These genes (fbln7 = LOC103040604; cyp24a1 = LOC103030576; spdef = LOC103037006; fam163b = LOC103027628) were relabeled with the gene names from the surface fish 2.0 genome annotation in the figures to maintain consistency.

## ACKNOWLEDGEMENTS

The authors thank all members of the Kowalko lab for help with fish care. We would like to acknowledge bioinformatics and sequencing support from the UH-Sequencing & Gene Editing Core. This study was supported by National Institutes of Health grant R15HD009022 and R35GM138345 to Johanna Kowalko.

## Reference

1. Courtier-Orgogozo, V., Arnoult, L., Prigent, S.R., Wiltgen, S., and Martin, A. (2020). Gephebase, a database of genotype-phenotype relationships for natural and domesticated variation in Eukaryotes. Nucleic Acids Res. 48, D696–D703.

2. Peichel, C.L., and Marques, D.A. (2017). The genetic and molecular architecture of phenotypic diversity in sticklebacks. Philos. Trans. R. Soc. Lond. B Biol. Sci. 372, 20150486.

3. Bendesky, A., Kwon, Y.-M., Lassance, J.-M., Lewarch, C.L., Yao, S., Peterson, B.K., He, M.X., Dulac, C., and Hoekstra, H.E. (2017). The genetic basis of parental care evolution in monogamous mice. Nature 544, 434–439.

4. Coughlan, J.M., Brown, M.W., and Willis, J.H. (2021). The genetic architecture and evolution of life-history divergence among perennials in the Mimulus guttatus species complex. Proc. Biol. Sci. 288, 20210077.

5. Herman, A., Brandvain, Y., Weagley, J., Jeffery, W.R., Keene, A.C., Kono, T.J.Y., Bilandžija, H., Borowsky, R., Espinasa, L., O’Quin, K., et al. (2018). The role of gene flow in rapid and repeated evolution of cave-related traits in Mexican tetra, Astyanax mexicanus. Mol. Ecol. 27, 4397–4416.

6. Garduño-Sánchez, M., Hernández-Lozano, J., Moran, R.L., Miranda-Gamboa, R., Gross, J.B., Rohner, N., Elliott, W.R., Miller, J., Lozano-Vilano, L., McGaugh, S.E., et al. (2023). Phylogeographic relationships and morphological evolution between cave and surface Astyanax mexicanus populations (De Filippi 1853) (Actinopterygii, Characidae). Mol. Ecol. 32, 5626–5644.

7. Jeffery, W.R. (2020). Astyanax surface and cave fish morphs. Evodevo 11, 14.

8. Kowalko, J. (2020). Utilizing the blind cavefish Astyanax mexicanus to understand the genetic basis of behavioral evolution. J. Exp. Biol. 223. 10.1242/jeb.208835.

9. Protas, M.E., Hersey, C., Kochanek, D., Zhou, Y., Wilkens, H., Jeffery, W.R., Zon, L.I., Borowsky, R., and Tabin, C.J. (2006). Genetic analysis of cavefish reveals molecular convergence in the evolution of albinism. Nat. Genet. 38, 107–111.

10. Protas, M., Conrad, M., Gross, J.B., Tabin, C., and Borowsky, R. (2007). Regressive Evolution in the Mexican Cave Tetra, Astyanax mexicanus. Curr. Biol. 17, 452–454.

11. Protas, M., Tabansky, I., Conrad, M., Gross, J.B., Vidal, O., Tabin, C.J., and Borowsky, R. (2008). Multi-trait evolution in a cave fish, Astyanax mexicanus. Evol. Dev. 10, 196–209.

12. Stahl, B.A., and Gross, J.B. (2015). Alterations in Mc1r gene expression are associated with regressive pigmentation in Astyanax cavefish. Dev. Genes Evol. 225, 367–375.

13. O’Quin, K.E., Yoshizawa, M., Doshi, P., and Jeffery, W.R. (2013). Quantitative Genetic Analysis of Retinal Degeneration in the Blind Cavefish Astyanax mexicanus. PLoS One 8, e57281.

14. Yoshizawa, M., Yamamoto, Y., O’Quin, K.E., and Jeffery, W.R. (2012). Evolution of an adaptive behavior and its sensory receptors promotes eye regression in blind cavefish. BMC Biol. 10, 108.

15. Kowalko, J.E., Rohner, N., Linden, T.A., Rompani, S.B., Warren, W.C., Borowsky, R., Tabin, C.J., Jeffery, W.R., and Yoshizawa, M. (2013). Convergence in feeding posture occurs through different genetic loci in independently evolved cave populations of Astyanax mexicanus. Proc. Natl. Acad. Sci. U. S. A. 110, 16933–16938.

16. Kowalko, J.E., Rohner, N., Rompani, S.B., Peterson, B.K., Linden, T.A., Yoshizawa, M., Kay, E.H., Weber, J., Hoekstra, H.E., Jeffery, W.R., et al. (2013). Loss of schooling behavior in cavefish through sight-dependent and sight-independent mechanisms. Curr. Biol. 23, 1874–1883.

17. Yoshizawa, M., Robinson, B.G., Duboué, E.R., Masek, P., Jaggard, J.B., O’Quin, K.E., Borowsky, R.L., Jeffery, W.R., and Keene, A.C. (2015). Distinct genetic architecture underlies the emergence of sleep loss and prey-seeking behavior in the Mexican cavefish. BMC Biol. 13, 15.

18. O’Quin, K.E., Doshi, P., Lyon, A., Hoenemeyer, E., Yoshizawa, M., and Jeffery, W.R. (2015). Complex evolutionary and genetic patterns characterize the loss of scleral ossification in the blind cavefish Astyanax mexicanus. PLoS One 10, e0142208.

19. Carlson, B.M., Klingler, I.B., Meyer, B.J., and Gross, J.B. (2018). Genetic analysis reveals candidate genes for activity QTL in the blind Mexican tetra, Astyanax mexicanus. PeerJ 6, e5189.

20. Riddle, M.R., Aspiras, A.C., Damen, F., Hutchinson, J.N., Chinnapen, D.J.-F., Tabin, J., and Tabin, C.J. (2020). Genetic architecture underlying changes in carotenoid accumulation during the evolution of the blind Mexican cavefish, Astyanax mexicanus. J. Exp. Zool. B Mol. Dev. Evol. 334, 405–422.

21. Riddle, M.R., Aspiras, A., Damen, F., McGaugh, S., Tabin, J.A., and Tabin, C.J. (2021). Genetic mapping of metabolic traits in the blind Mexican cavefish reveals sex-dependent quantitative trait loci associated with cave adaptation. BMC Ecol. Evol. 21, 94.

22. Powers, A.K., Hyacinthe, C., Riddle, M.R., Kim, Y.K., Amaismeier, A., Thiel, K., Martineau, B., Ferrante, E., Moran, R.L., McGaugh, S.E., et al. (2023). Genetic mapping of craniofacial traits in the Mexican tetra reveals loci associated with bite differences between cave and surface fish. BMC Ecol. Evol. 23, 41.

23. Powers, A.K., Amaismeier, A., Thiel, K., Anyonge, W., McGaugh, S.E., Boggs, T.E., Tabin, C.J., and Gross, J.B. (2025). Genetic mapping of orofacial traits reveals a single genomic region associated with differences in multiple parameters of jaw size between Astyanax mexicanus surface and cavefish. Evol. Dev., e70003.

24. McGaugh, S.E., Gross, J.B., Aken, B., Blin, M., Borowsky, R., Chalopin, D., Hinaux, H., Jeffery, W.R., Keene, A., Ma, L., et al. (2014). The cavefish genome reveals candidate genes for eye loss. Nat. Commun. 5, 5307.

25. Warren, W.C., Boggs, T.E., Borowsky, R., Carlson, B.M., Ferrufino, E., Gross, J.B., Hillier, L., Hu, Z., Keene, A.C., Kenzior, A., et al. (2021). A chromosome-level genome of Astyanax mexicanus surface fish for comparing population-specific genetic differences contributing to trait evolution. Nat. Commun. 12, 1447.

26. Wiese, J., Richards, E., Kowalko, J.E., and McGaugh, S.E. (2024). Quantitative trait loci concentrate in specific regions of the Mexican cavefish genome and reveal key candidate genes for cave-associated evolution. J. Hered. 10.1093/jhered/esae040.

27. Klaassen, H., Wang, Y., Adamski, K., Rohner, N., and Kowalko, J.E. (2018). CRISPR mutagenesis confirms the role of oca2 in melanin pigmentation in Astyanax mexicanus. Dev. Biol. 441, 313–318.

28. Ma, L., Gore, A.V., Castranova, D., Shi, J., Ng, M., Tomins, K.A., van der Weele, C.M., Weinstein, B.M., and Jeffery, W.R. (2020). A hypomorphic cystathionine ß-synthase gene contributes to cavefish eye loss by disrupting optic vasculature. Nat. Commun. 11, 2772.

29. Shennard, D., Sifuentes-Romero, I., Ambosie, R., Abdelaziz, J., Duboue, E.R., and Kowalko, J.E. (2025). The rx3 gene contributes to the evolution of eye loss in the cavefish Astyanax mexicanus. Evol. Dev. 27, e70011.

30. Sifuentes-Romero, I., Aviles, A.M., Carter, J.L., Chan-Pong, A., Clarke, A., Crotty, P., Engstrom, D., Meka, P., Perez, A., Perez, R., et al. (2023). Trait loss in evolution: What cavefish have taught us about mechanisms underlying eye regression. Integr. Comp. Biol. 63, 393–406.

31. Jeffery, W.R., Strickler, A.G., and Yamamoto, Y. (2003). To see or not to see: evolution of eye degeneration in mexican blind cavefish. Integr. Comp. Biol. 43, 531–541.

32. Krishnan, J., and Rohner, N. (2017). Cavefish and the basis for eye loss. Philos. Trans. R. Soc. Lond. B Biol. Sci. 372. 10.1098/rstb.2015.0487.

33. Gore, A.V., Tomins, K.A., Iben, J., Ma, L., Castranova, D., Davis, A.E., Parkhurst, A., Jeffery, W.R., and Weinstein, B.M. (2018-7). An epigenetic mechanism for cavefish eye degeneration. Nat Ecol Evol 2, 1155–1160.

34. Moran, R.L., Richards, E.J., Ornelas-García, C.P., Gross, J.B., Donny, A., Wiese, J., Keene, A.C., Kowalko, J.E., Rohner, N., and McGaugh, S.E. (2023). Selection-driven trait loss in independently evolved cavefish populations. Nat. Commun. 14, 2557.

35. Clark, D.T. (1981). Visual responses in developing zebrafish (*Brachydanio rerio*).

36. Neuhauss, S.C.F. (2003). Behavioral genetic approaches to visual system development and function in zebrafish. J. Neurobiol. 54, 148–160.

37. Neuhauss, S.C.F., Biehlmaier, O., Seeliger, M.W., Das, T., Kohler, K., Harris, W.A., and Baier, H. (1999). Genetic Disorders of Vision Revealed by a Behavioral Screen of 400 Essential Loci in Zebrafish. J. Neurosci. 19, 8603–8615.

38. Chuang, J.C., Mathers, P.H., and Raymond, P.A. (1999). Expression of three Rx homeobox genes in embryonic and adult zebrafish. Mech. Dev. 84, 195–198.

39. Deschet, K., Bourrat, F., Ristoratore, F., Chourrout, D., and Joly, J.S. (1999). Expression of the medaka (Oryzias latipes) Ol-Rx3 paired-like gene in two diencephalic derivatives, the eye and the hypothalamus. Mech. Dev. 83, 179–182.

40. Mathers, P.H., Grinberg, A., Mahon, K.A., and Jamrich, M. (1997). The Rx homeobox gene is essential for vertebrate eye development. Nature 387, 603–607.

41. Muranishi, Y., Terada, K., and Furukawa, T. (2012). An essential role for Rax in retina and neuroendocrine system development. Dev. Growth Differ. 54, 341–348.

42. Kennedy, B.N., Stearns, G.W., Smyth, V.A., Ramamurthy, V., van Eeden, F., Ankoudinova, I., Raible, D., Hurley, J.B., and Brockerhoff, S.E. (2004). Zebrafish *rx3* and *mab21l2* are required during eye morphogenesis. Dev. Biol. 270, 336–349.

43. Loosli, F., Staub, W., Finger-Baier, K.C., Ober, E.A., Verkade, H., Wittbrodt, J., and Baier, H. (2003). Loss of eyes in zebrafish caused by mutation of chokh/rx3. EMBO Rep. 4, 894–899.

44. Winkler, S., Loosli, F., Henrich, T., Wakamatsu, Y., and Wittbrodt, J. (2000). The conditional medaka mutation eyeless uncouples patterning and morphogenesis of the eye. Development 127, 1911–1919.

45. Chakraborty, P., Dash, S.P., and Sarangi, P.P. (2020). The role of adhesion protein Fibulin7 in development and diseases. Mol. Med. 26, 47.

46. Ikeuchi, T., de Vega, S., Forcinito, P., Doyle, A.D., Amaral, J., Rodriguez, I.R., Arikawa-Hirasawa, E., and Yamada, Y. (2018). Extracellular protein fibulin-7 and its C-terminal fragment have in vivo antiangiogenic activity. Sci. Rep. 8, 17654.

47. Culver, D.C., and Pipan, T. (2019). The Biology of Caves and Other Subterranean Habitats. 10.1093/oso/9780198820765.001.0001.

48. Pottin, K., Hinaux, H., and Rétaux, S. (2011). Restoring eye size in Astyanax mexicanus blind cavefish embryos through modulation of the Shh and Fgf8 forebrain organising centres. Development 138, 2467–2476.

49. Alunni, A., Menuet, A., Candal, E., Pénigault, J.-B., Jeffery, W.R., and Rétaux, S. (2007). Developmental mechanisms for retinal degeneration in the blind cavefish Astyanax mexicanus. J. Comp. Neurol. 505, 221–233.

50. Jeffery, W.R., and Martasian, D.P. (1998). Evolution of Eye Regression in the Cavefish Astyanax: Apoptosis and the Pax-6 Gene1. Am. Zool. 38, 685–696.

51. Strickler, A.G., Byerly, M.S., and Jeffery, W.R. (2007). Lens gene expression analysis reveals downregulation of the anti-apoptotic chaperone alphaA-crystallin during cavefish eye degeneration. Dev. Genes Evol. 217, 771–782.

52. Yamamoto, Y., Stock, D.W., and Jeffery, W.R. (2004). Hedgehog signalling controls eye degeneration in blind cavefish. Nature 431, 844–847.

53. Yamamoto, Y., and Jeffery, W.R. (2000). Central role for the lens in cave fish eye degeneration. Science 289, 631–633.

54. Sifuentes-Romero, I., Ferrufino, E., Thakur, S., Laboissonniere, L.A., Solomon, M., Smith, C.L., Keene, A.C., Trimarchi, J.M., and Kowalko, J.E. (2020). Repeated evolution of eye loss in Mexican cavefish: Evidence of similar developmental mechanisms in independently evolved populations. J. Exp. Zool. B Mol. Dev. Evol. 334, 423–437.

55. Leclercq, J., Torres-Paz, J., Policarpo, M., Agnès, F., and Rétaux, S. (2024). Evolution of the regulation of developmental gene expression in blind Mexican cavefish. Development 151, dev202610.

56. Policarpo, M., Legendre, L., Germon, I., Lafargeas, P., Espinasa, L., Rétaux, S., and Casane, D. (2024). The nature and distribution of putative non-functional alleles suggest only two independent events at the origins of Astyanax mexicanus cavefish populations. BMC Ecol. Evol. 24, 41.

57. Roback, E.Y., Ferrufino, E., Moran, R.L., Shennard, D., Mulliniks, C., Gallop, J., Weagley, J., Miller, J., Fily, Y., Ornelas-García, C.P., et al. (2025). Population genomics of premature termination codons in cavefish with substantial trait loss. Mol. Biol. Evol. 42, msaf012.

58. Raymond, S.M., and Jackson, I.J. (1995). The retinal pigmented epithelium is required for development and maintenance of the mouse neural retina. Curr. Biol. 5, 1286–1295.

59. Ma, L., Ng, M., van der Weele, C.M., Yoshizawa, M., and Jeffery, W.R. (2020). Dual roles of the retinal pigment epithelium and lens in cavefish eye degeneration. J. Exp. Zool. B Mol. Dev. Evol. 334, 438–449.

60. Li, M., Zhao, L., Page-McCaw, P.S., and Chen, W. (2016). Zebrafish genome engineering using the CRISPR-Cas9 system. Trends Genet. 32, 815–827.

61. Hall, B., Cho, A., Limaye, A., Cho, K., Khillan, J., and Kulkarni, A.B. (2018). Genome editing in mice using CRISPR/Cas9 technology. Curr. Protoc. Cell Biol. 81, e57.

62. Gratz, S.J., Rubinstein, C.D., Harrison, M.M., Wildonger, J., and O’Connor-Giles, K.M. (2015). CRISPR-Cas9 genome editing in Drosophila. Curr. Protoc. Mol. Biol. 111, 31.2.1-31.2.20.

63. Lu, F., Leach, L.L., and Gross, J.M. (2023). A CRISPR-Cas9-mediated F0 screen to identify pro-regenerative genes in the zebrafish retinal pigment epithelium. Sci. Rep. 13, 3142.

64. Klatt Shaw, D., and Mokalled, M.H. (2021). Efficient CRISPR/Cas9 mutagenesis for neurobehavioral screening in adult zebrafish. G3 (Bethesda) 11, jkab089.

65. Lin, W., Shi, Y., Tian, J., Liu, X., Weng, F., and Wu, Z. (2025). Kdm7aa orchestrates an immunomodulatory cardiomyocyte program to enable zebrafish heart regeneration. Int. J. Mol. Sci. 26, 10044.

66. Gilles, A.F., Schinko, J.B., and Averof, M. (2015). Efficient CRISPR-mediated gene targeting and transgene replacement in the beetle Tribolium castaneum. Development 142, 2832–2839.

67. Klingler, M., and Bucher, G. (2022). The red flour beetle T. castaneum: elaborate genetic toolkit and unbiased large scale RNAi screening to study insect biology and evolution. Evodevo 13, 14.

68. Wucherpfennig, J.I., Miller, C.T., and Kingsley, D.M. (2019). Efficient CRISPR-Cas9 editing of major evolutionary loci in sticklebacks. Evol. Ecol. Res. 20, 107–132.

69. Martin, A., Serano, J.M., Jarvis, E., Bruce, H.S., Wang, J., Ray, S., Barker, C.A., O’Connell, L.C., and Patel, N.H. (2016). CRISPR/Cas9 Mutagenesis reveals versatile roles of Hox genes in crustacean limb specification and evolution. Curr. Biol. 26, 14–26.

70. O’Gorman, M., Thakur, S., Imrie, G., Moran, R.L., Choy, S., Sifuentes-Romero, I., Bilandžija, H., Renner, K.J., Duboué, E., Rohner, N., et al. (2021). Pleiotropic function of the oca2 gene underlies the evolution of sleep loss and albinism in cavefish. Curr. Biol. 31, 3694–3701.e4.

71. Choy, S., Thakur, S., Polyakov, E., Abdelaziz, J., Lloyd, E., Enriquez, M., Jayan, N., Mensinger, A., Fily, Y., McGaugh, S., et al. (2025). Mutations in the albinism gene oca2 alter vision-dependent prey capture behavior in the Mexican tetra. J. Exp. Biol. 228, jeb249881.

72. Mack, K.L., Jaggard, J.B., Persons, J.L., Roback, E.Y., Passow, C.N., Stanhope, B.A., Ferrufino, E., Tsuchiya, D., Smith, S.E., Slaughter, B.D., et al. (2021). Repeated evolution of circadian clock dysregulation in cavefish populations. PLoS Genet. 17, e1009642.

73. Perera, P.P., Guerra, D.P., and Riddle, M.R. (2023). The Mexican Tetra, Astyanax mexicanus, as a Model System in Cell and Developmental Biology. Annu. Rev. Cell Dev. Biol. 39, 23–44.

74. Blin, M., Tine, E., Meister, L., Elipot, Y., Bibliowicz, J., Espinasa, L., and Rétaux, S. (2018). Developmental evolution and developmental plasticity of the olfactory epithelium and olfactory skills in Mexican cavefish. Dev. Biol. 441, 242–251.

75. Fernandes, V.F.L., Macaspac, C., Lu, L., and Yoshizawa, M. (2018). Evolution of the developmental plasticity and a coupling between left mechanosensory neuromasts and an adaptive foraging behavior. Dev. Biol. 441, 262–271.

76. Franz-Odendaal, T.A., and Hall, B.K. (2006). Modularity and sense organs in the blind cavefish, Astyanax mexicanus. Evol. Dev. 8, 94–100.

77. Jeffery, W.R. (2005). Adaptive Evolution of Eye Degeneration in the Mexican Blind Cavefish. J. Hered. 96, 185–196.

78. Jeffery, W.R. (2009). Evolution and development in the cavefish Astyanax. Curr. Top. Dev. Biol. 86, 191–221.

79. Maldonado, E., Rangel-Huerta, E., Rodriguez-Salazar, E., Pereida-Jaramillo, E., and Martínez-Torres, A. (2020). Subterranean life: Behavior, metabolic, and some other adaptations of Astyanax cavefish. J. Exp. Zool. B Mol. Dev. Evol. 334, 463–473.

80. Menuet, A., Alunni, A., Joly, J.-S., Jeffery, W.R., and Rétaux, S. (2007). Expanded expression of Sonic Hedgehog in Astyanax cavefish:multiple consequences on forebrain development and evolution. Development 134, 845–855.

81. Duboué, E.R., Keene, A.C., and Borowsky, R.L. (2011). Evolutionary Convergence on Sleep Loss in Cavefish Populations. Curr. Biol. 21, 671–676.

82. Elipot, Y., Hinaux, H., Callebert, J., and Rétaux, S. (2013). Evolutionary Shift from Fighting to Foraging in Blind Cavefish through Changes in the Serotonin Network. Curr. Biol. 23, 1–10.

83. Patch, A., Paz, A., Holt, K.J., Duboué, E.R., Keene, A.C., Kowalko, J.E., and Fily, Y. (2022). Kinematic analysis of social interactions deconstructs the evolved loss of schooling behavior in cavefish. PLoS One 17, e0265894.

84. Rodriguez-Morales, R., Gonzalez-Lerma, P., Yuiska, A., Han, J.H., Guerra, Y., Crisostomo, L., Keene, A.C., Duboue, E.R., and Kowalko, J.E. (2022). Convergence on reduced aggression through shared behavioral traits in multiple populations of Astyanax mexicanus. BMC Ecology and Evolution 22, 116.

85. Warren, W.C., Rice, E.S., Maggs, X., Roback, E., Keene, A., Martin, F., Ogeh, D., Haggerty, L., Carroll, R.A., McGaugh, S., et al. (2023). Astyanax mexicanus surface and cavefish chromosome-scale assemblies for trait variation discovery. 10.1101/2023.11.16.567450.

86. Gross, J.B., Furterer, A., Carlson, B.M., and Stahl, B.A. (2013). An Integrated Transcriptome-Wide Analysis of Cave and Surface Dwelling Astyanax mexicanus. PLoS One 8, e55659.

87. Krug, J., Perner, B., Albertz, C., Mörl, H., Hopfenmüller, V.L., and Englert, C. (2023). Generation of a transparent killifish line through multiplex CRISPR/Cas9mediated gene inactivation. Elife 12. 10.7554/eLife.81549.

88. Kim, Y., Nam, H.G., and Valenzano, D.R. (2016). The short-lived African turquoise killifish: an emerging experimental model for ageing. Dis. Model. Mech. 9, 115–129.

89. Clark, B., Kuwalekar, M., Fischer, B., Woltering, J., Biran, J., Juntti, S., Kratochwil, C.F., Santos, M.E., and Almeida, M.V. (2023). Genome editing in East African cichlids and tilapias: state-of-the-art and future directions. Open Biol. 13, 230257.

90. Kern, A.D., and Schrider, D.R. (2018). DiploS/HIC: An updated approach to classifying selective sweeps. G3 (Bethesda) 8, 1959–1970.

91. Montague, T.G., Cruz, J.M., Gagnon, J.A., Church, G.M., and Valen, E. (2014). CHOPCHOP: a CRISPR/Cas9 and TALEN web tool for genome editing. Nucleic Acids Res. 42, W401–7.

92. Stahl, B.A., Jaggard, J.B., Chin, J.S.R., Kowalko, J.E., Keene, A.C., and Duboué, E.R. (2019). Manipulation of gene function in Mexican cavefish. J. Vis. Exp., e59093.

93. Kroll, F., Powell, G.T., Ghosh, M., Gestri, G., Antinucci, P., Hearn, T.J., Tunbak, H., Lim, S., Dennis, H.W., Fernandez, J.M., et al. (2021). A simple and effective F0 knockout method for rapid screening of behaviour and other complex phenotypes. Elife 10. 10.7554/eLife.59683.

94. Schneider, C.A., Rasband, W.S., and Eliceiri, K.W. (2012-7). NIH Image to ImageJ: 25 years of Image Analysis. Nat. Methods 9, 671–675.

95. Schindelin, J., Arganda-Carreras, I., Frise, E., Kaynig, V., Longair, M., Pietzsch, T., Preibisch, S., Rueden, C., Saalfeld, S., Schmid, B., et al. (2012). Fiji: an open-source platform for biological-image analysis. Nat. Methods 9, 676–682.

96. Kowalko, J.E., Ma, L., and Jeffery, W.R. (2016). Genome editing in Astyanax mexicanus using transcription activator-like effector nucleases (TALENs). J. Vis. Exp. 10.3791/54113.

97. Sifuentes-Romero, I., Ferrufino, E., and Kowalko, J.E. (2023). Application of CRISPR-Cas9 for Functional Analysis in A. mexicanus. In Neuromethods Neuromethods. (Springer US), pp. 193–220.

98. Clement, K., Rees, H., Canver, M.C., Gehrke, J.M., Farouni, R., Hsu, J.Y., Cole, M.A., Liu, D.R., Joung, J.K., Bauer, D.E., et al. (2019). CRISPResso2 provides accurate and rapid genome editing sequence analysis. Nat. Biotechnol. 37, 224–226.

99. Larsson, A. (2014). AliView: a fast and lightweight alignment viewer and editor for large datasets. Bioinformatics 30, 3276–3278.

100. Blanchette, M., Kent, W.J., Riemer, C., Elnitski, L., Smit, A.F.A., Roskin, K.M., Baertsch, R., Rosenbloom, K., Clawson, H., Green, E.D., et al. (2004). Aligning multiple genomic sequences with the threaded blockset aligner. Genome Res. 14, 708–715.

101. Perez, G., Barber, G.P., Benet-Pages, A., Casper, J., Clawson, H., Diekhans, M., Fischer, C., Gonzalez, J.N., Hinrichs, A.S., Lee, C.M., et al. (2025). The UCSC Genome Browser database: 2025 update. Nucleic Acids Res. 53, D1243–D1249.

102. Morales, J., Pujar, S., Loveland, J.E., Astashyn, A., Bennett, R., Berry, A., Cox, E., Davidson, C., Ermolaeva, O., Farrell, C.M., et al. (2022). A joint NCBI and EMBL-EBI transcript set for clinical genomics and research. Nature 604, 310–315.

103. Edgar, R.C. (2004). MUSCLE: multiple sequence alignment with high accuracy and high throughput. Nucleic Acids Res. 32, 1792–1797.

104. Gouy, M., Guindon, S., and Gascuel, O. (2010). SeaView version 4: A multiplatform graphical user interface for sequence alignment and phylogenetic tree building. Mol. Biol. Evol. 27, 221–224.

105. Zheng, G.X.Y., Terry, J.M., Belgrader, P., Ryvkin, P., Bent, Z.W., Wilson, R., Ziraldo, S.B., Wheeler, T.D., McDermott, G.P., Zhu, J., et al. (2017). Massively parallel digital transcriptional profiling of single cells. Nat. Commun. 8, 14049.

106. Hao, Y., Stuart, T., Kowalski, M.H., Choudhary, S., Hoffman, P., Hartman, A., Srivastava, A., Molla, G., Madad, S., Fernandez-Granda, C., et al. (2024). Dictionary learning for integrative, multimodal and scalable single-cell analysis. Nat. Biotechnol. 42, 293–304.

107. McGinnis, C.S., Murrow, L.M., and Gartner, Z.J. (2019). DoubletFinder: Doublet Detection in Single-Cell RNA Sequencing Data Using Artificial Nearest Neighbors. Cell Syst 8, 329–337.e4.

