## Supplemental Data for "CRISPR screening reveals genetic regulators associated with the evolution of eye degeneration"

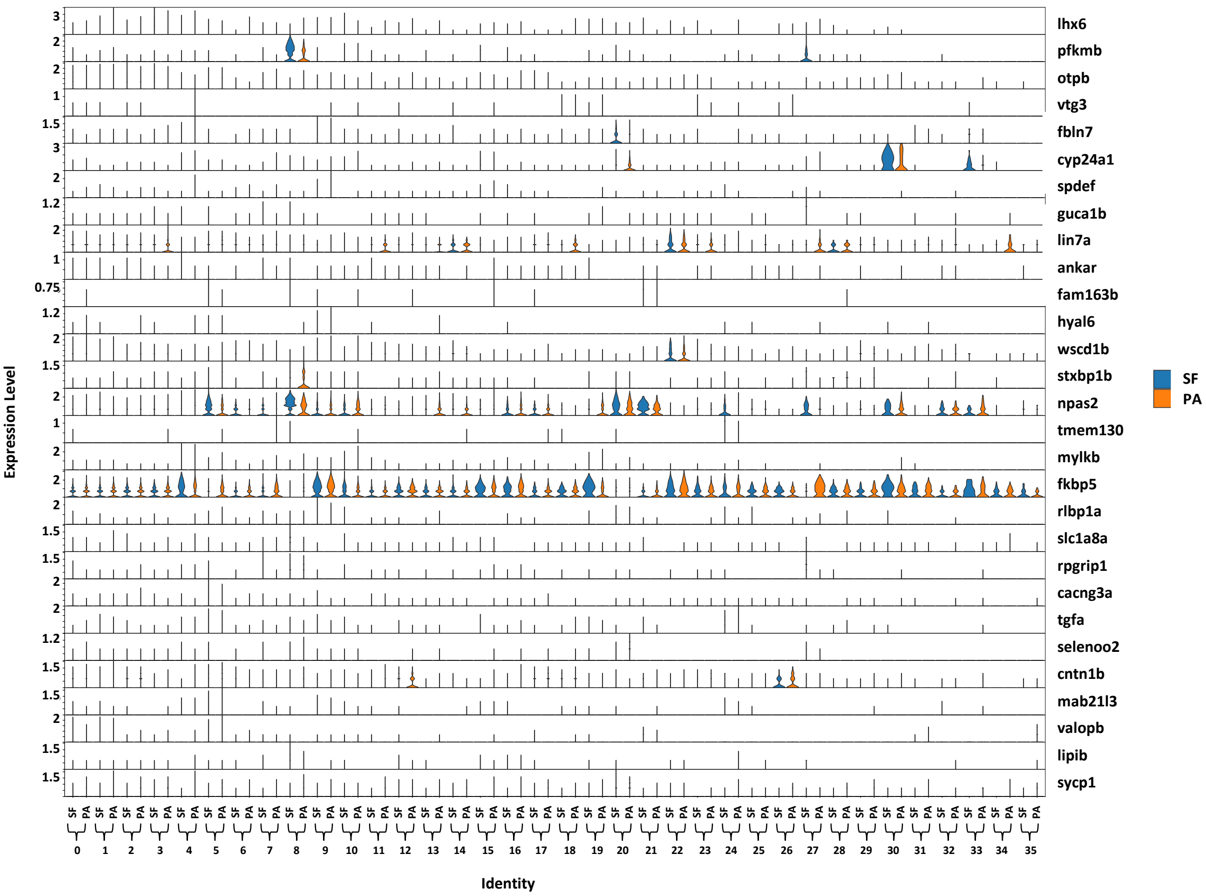


**Supplemental Figure 1 – Expression of eye candidate genes in each cell cluster in surface and cavefish.**
Expression levels of each eye candidate gene are shown for surface fish (SF, blue) and Pachón cavefish (CF, pink) in each cell cluster derived from snRNA-sequencing. Differential expression analysis was performed to determine which genes were differentially expressed within each cluster between surface and cave fish (see Supplemental Table 1).

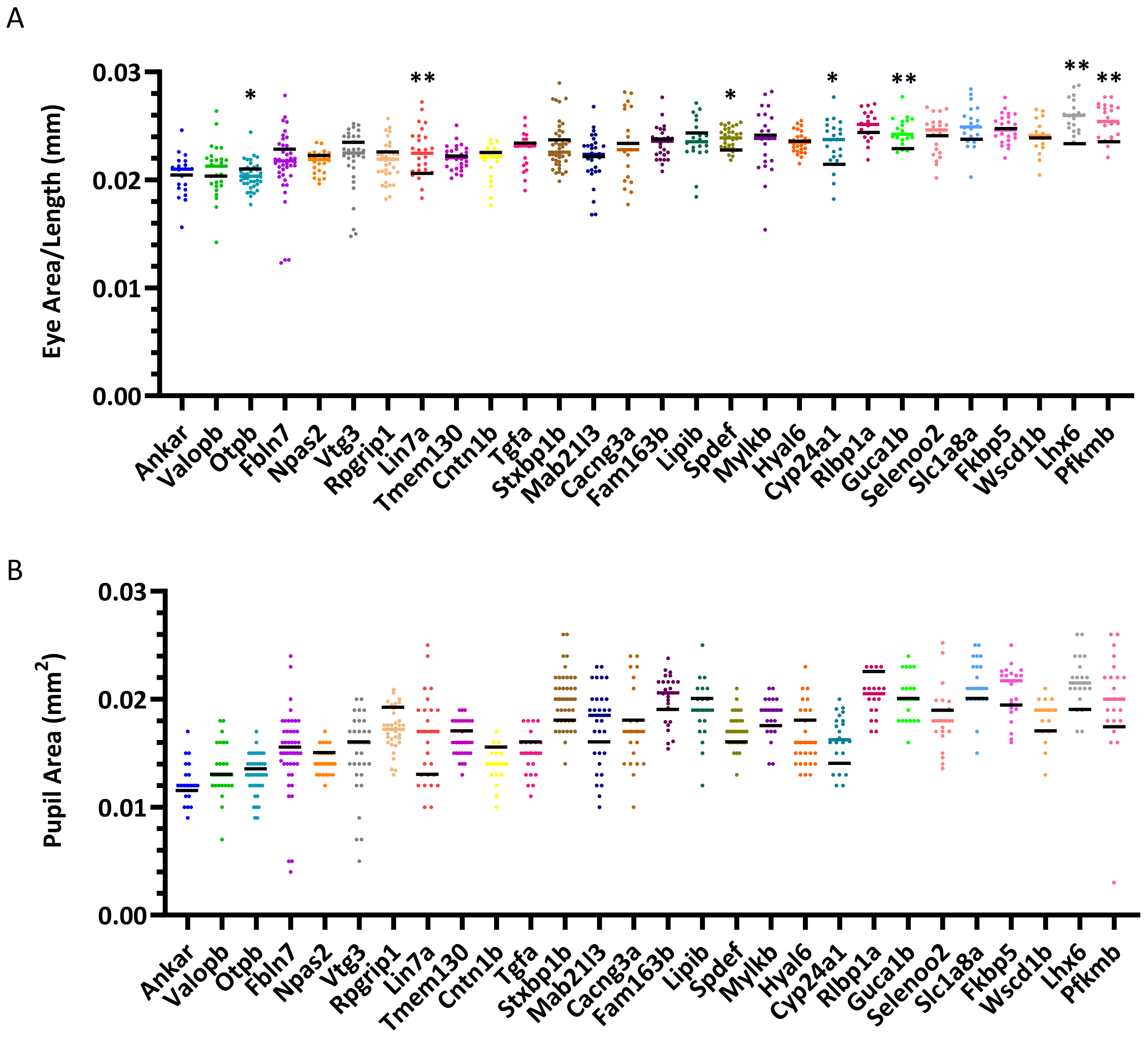


**Supplemental Figure 2 – Right eye and pupil size in crispant fish compared to wild-type siblings**

Right eye area and pupil area in crispant fish compared to wild-type siblings. Each data point represents 1 crispant fish and the colored bar is the median of the crispant fish. The black bar indicates the median of the wild-type siblings. All data corresponds to left eye and pupil data reported in Figure 4. All individual values are reported in Supplemental table 3 and statistics are reported in Supplemental Table 4.

A

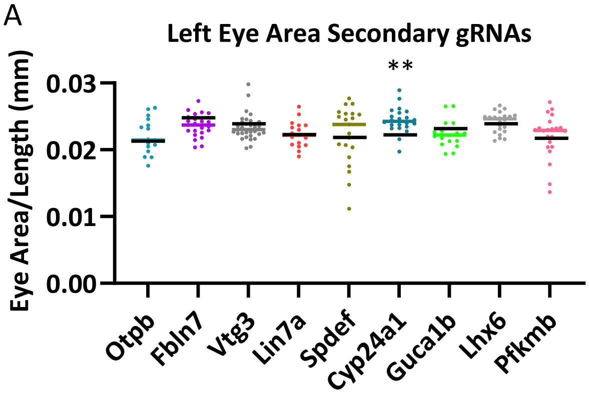


B

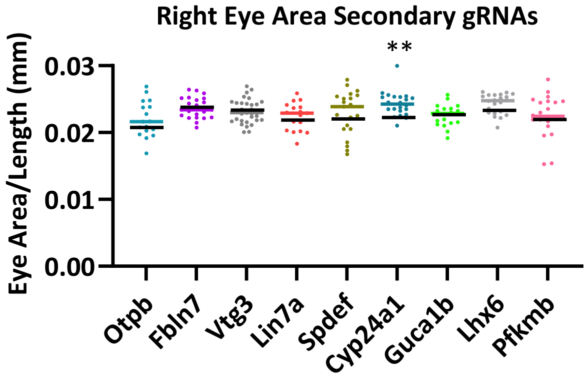


**Supplemental Figure 3 – Genes investigated with secondary gRNA and scramble control.**
All genes that resulted in differences in eye size when targeted in the primary screen were further investigated utilizing a secondary gRNA targeting the gene of interest and a scramble control. Each colored data point represents a single individual crispant fish and the colored bar indicates the median of the crispant fish. The black bar indicates the median of the scramble injected control siblings. Left eye area (A) and right eye area (B) from the secondary screen are shown. All individual values are reported in Supplemental table 3.

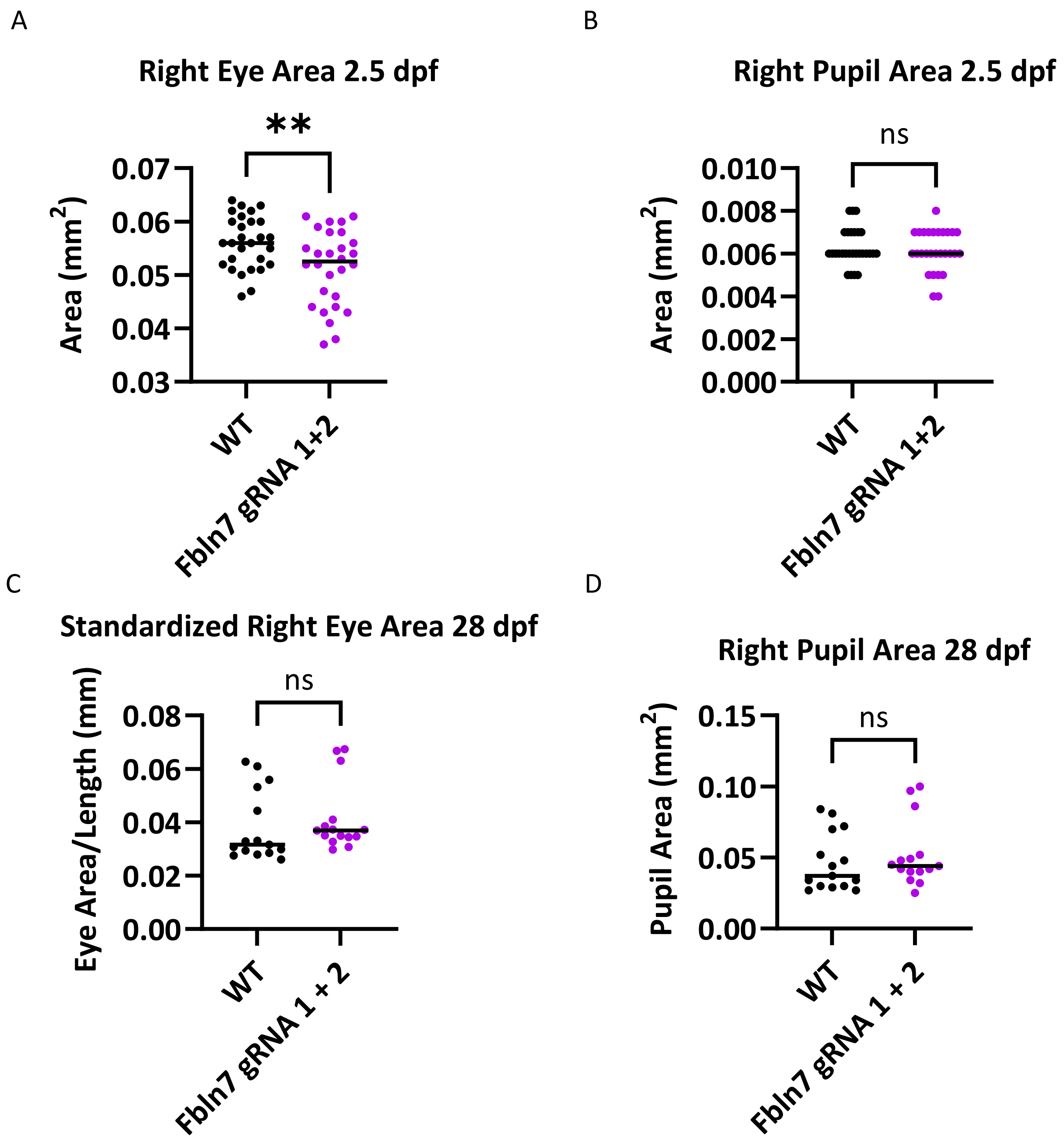


**Supplemental Figure 4 – Right eye area in *fbln7* gRNA 1 + 2 injected fish.**
Right eye (a,c) and pupil area (b,d) for *fbln7* gRNA 1 + 2 crispant injected fish and wild-type siblings at 2.5 dpf (a,b) and 28 dpf (c,d) Corresponds to left eye and pupil data reported in Figure 5.

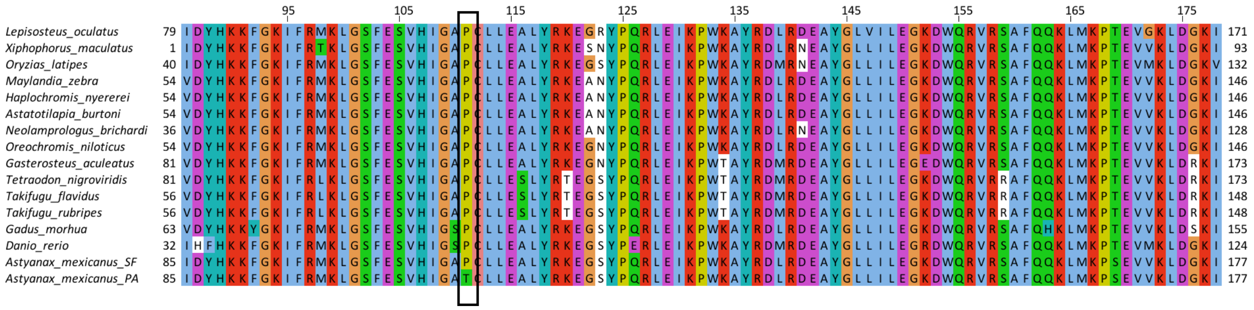


**Supplemental Figure 5 - Alignment of Cyp24a1 protein sequence across species showing conserved coding change in Pachón cavefish.** Part of the protein coding sequence alignment. Numbers on left and right of alignments indicate amino acid position within species. Box shows position at which Pachón cavefish exhibit a coding change relative to other species.

### Supplemental Tables

**Supplemental Table 1 – Differential gene expression analysis for eye candidate genes.** The snRNA-sequencing data was evaluated to determine if the 29 candidate genes were differentially expressed in surface fish relative to Pachón cavefish in each cell cluster. Each tab includes differential expression data for each cluster. Positive values in the avg_log2FC column indicate higher expression in surface fish, and negative numbers indicate higher expression in cavefish.

**Supplemental Table 2 – Sequences for gRNAs and primers**
gRNAs to target each of the eye candidate genes were designed using ChopChop. Primers were designed to amplify the target sequence.

**Supplemental Table 3 – Phenotyping data for all figures**
Individual values for all phenotyping data across Figures 2,3,4,5 and Supplemental figures 2,3,4.

**Supplemental Table 4 – Statistics for eye candidate genes crispant screen**
All statistical analyses for Figures 3, 4 and Supplemental Figure 2, 3.

**Supplemental Table 5 – Amplicon sequencing results.** Results in percentage altered alleles for all gRNAs per embryo.

**Supplemental Table 6 – Analysis of coding changes in Pachon cavefish.**

**Supplemental Table 7- Enriched genes for snRNA-seq clusters.**
